# Evolution and molecular basis of a novel allosteric property of crocodilian hemoglobin

**DOI:** 10.1101/2022.07.18.500494

**Authors:** Chandrasekhar Natarajan, Anthony V. Signore, Naim M. Bautista, Federico G. Hoffmann, Jeremy R. H. Tame, Angela Fago, Jay F. Storz

## Abstract

Understanding the evolution of novel protein functions requires data on the mechanistic effects of causative mutations and the extent of coupling between the gain of new function and loss of ancestral function. Here, we use ancestral protein resurrection and directed mutagenesis to dissect the molecular basis of a novel mode of allosteric regulation in crocodilian hemoglobin. We discovered that regulation of Hb-O_2_ affinity via allosteric binding of bicarbonate ions (a biochemical adaptation unique to crocodilians) and the concomitant loss of allosteric regulation by ATP-binding are attributable to non-overlapping sets of substitutions. Gain of bicarbonate-sensitivity involved direct effects of few replacements at key sites in combination with indirect effects of numerous replacements at structurally disparate sites. Due to the context-dependence of causative substitutions, the unique allosteric properties of crocodilian hemoglobin cannot be easily transplanted into divergent homologs of other species.

**Significance Statement:** The extraordinary breath-hold diving capacity of crocodilians is partly attributable to a unique mode of allosterically regulating hemoglobin-oxygenation in circulating red blood cells. We investigated the origin and mechanistic basis of this novel biochemical adaptation by performing directed mutagenesis experiments on resurrected ancestral crocodilian hemoglobins. Our results revealed that evolved changes in allosteric regulation involved the direct effect of few amino acid substitutions at key sites in combination with indirect effects of numerous other substitutions at structurally disparate sites. Such indirect interaction effects suggest that the evolution of novel protein functions may often depend on neutral mutations that produce no adaptive benefit when they first arise, but which contribute to a permissive background for subsequent function-altering mutations at other sites.

## Introduction

How do novel protein functions evolve in an incremental, step-by-step fashion, and to what extent is the gain of new function mechanistically coupled with the loss of ancestral function? These are fundamental questions in molecular evolution and they can be addressed with a protein-engineering approach that permits the identification and functional characterization of causative mutations. Here we report discoveries regarding the molecular basis of a key physiological innovation during vertebrate evolution. We used ancestral protein resurrection in conjunction with site-directed mutagenesis experiments to dissect the molecular basis of a unique mechanism for allosterically regulating the O_2_-affinity of crocodilian hemoglobin (Hb).

Crocodilians are able to remain submerged underwater for extraordinarily long periods of time – a physiological capacity that allows some species, such as the Nile crocodile (*Crocodylus niloticus*), to kill large, mammalian prey like wildebeest and zebras by dragging them underwater and drowning them. This capacity for breath-hold diving is made possible by the fact that the O_2_-affinity of crocodilian Hb is regulated by the inhibitory effects of bicarbonate ions (1-6). During submergence, a progressive increase in the red cell concentration of bicarbonate (derived from the hydration of CO_2_) reduces Hb-O_2_ affinity, thereby promoting O_2_ unloading to the cells of metabolizing tissues (5). This is a highly efficient physiological strategy since the circulatory O_2_ store can be almost fully depleted and the CO_2_ carrying capacity of the blood is maximized. As stated by Perutz (3): “It is surprising that the crocodilian hemoglobins’ simple and direct reciprocating action between oxygen and one of the end products of oxidative metabolism has not been adopted by other vertebrates”. The bicarbonate sensitivity of crocodilian Hb may also play an important role in tissue O_2_ delivery during the post-prandial ‘alkaline tide’, as the secretion of hydrochloric acid into the stomach (which aids the digestion of skeletal material) involves an anion exchange across the gut lining that reduces the blood concentration of chloride while increasing the blood pH and bicarbonate concentration (7-9).

The Hbs of jawed vertebrates are heterotetramers, consisting of two α-type subunits and two β-type subunits. Each of the four globin subunits contain a heme group with a ferrous (Fe^2+^) iron atom that reversibly binds a single O_2_ molecule. The Hb tetramer undergoes an oxygenation-linked transition in quaternary structure, whereby the two semi-rigid α_1_β_1_ and α_2_β_2_ dimers rotate relative to one another during the transition between the deoxy (low-affinity [T]) conformation and the oxy (high-affinity [R]) conformation (10). This oxygenation-linked shift in quaternary structure is central to the regulation of Hb-O_2_ affinity by allosteric cofactors (non-heme ligands such as hydrogen ions, chloride ions, and organic phosphates) that are present in the red blood cell. Allosteric cofactors reduce Hb-O_2_ affinity by preferentially binding to deoxyHb, thereby stabilizing the low-affinity T-state through the formation of additional hydrogen-bonds and salt bridges within and between subunits. Among vertebrate Hbs, crocodilian Hb is unique in that its O_2_-affinity is primarily modulated by bicarbonate ions rather than organic phosphates such as ATP. The origin and mechanistic basis of this unique mode of allosteric regulatory control has remained a mystery.

On the basis of structural modeling, Perutz et al. (2) hypothesized that the binding site for organic phosphates in the positively charged cleft between the β-chains was co-opted for allosteric bicarbonate-binding in crocodilian Hb, a shift that required a stereochemical reconfiguration of multiple residues. Perutz (3) suggested that the preferential binding of bicarbonate ions may have involved as few as three amino acid substitutions, two of which simultaneously eliminated residues directly involved in ATP-binding. According to this hypothesis, two additional amino acid substitutions (1 in the α-chain, 1 in the β-chain) would be required to completely abolish the ancestral sensitivity to ATP. These hypotheses were tested by engineering crocodile-specific mutations into recombinant human Hb (11). Contrary to Perutz’s predictions, the study by Komiyama et al. (11) demonstrated that transplanting bicarbonate sensitivity into human Hb required a total of 12 amino acid substitutions (7 in the α-chain, 5 in the β-chain), most of which were concentrated in the symmetrical α_1_β_2_ and α_2_β_1_ intersubunit contact surfaces.

Although the Komiyama *et al*. study provided important insights into possible structural mechanisms underlying bicarbonate-sensitivity, one problem with the interpretation of such ‘horizontal’ comparisons (swapping residues between homologous proteins of contemporary species) is that the focal mutations are introduced into a sequence context that may not be evolutionarily relevant. If mutations have context-dependent effects, then introducing crocodile-specific substitutions into human Hb may not recapitulate the functional effects of causative mutations on the genetic background in which they actually occurred during evolution (i.e., in the ancestor of modern crocodilians). Consequently, the particular substitutions required to transplant bicarbonate sensitivity into human Hb may be different from the substitutions that were responsible for the evolved bicarbonate sensitivity in the Hb of the crocodilian ancestor. An alternative ‘vertical’ approach is to reconstruct and resurrect ancestral proteins to test the effects of historical mutations on the genetic background in which they actually occurred during evolution (12-14). We used this approach to measure the functional effects of amino acid substitutions that occurred in the reconstructed ancestor of archosaurs, the group containing crocodilians and birds. Since bicarbonate-sensitivity is a property specific to the Hbs of crocodilians, it must have evolved in the line of descent leading from the archosaur ancestor, which existed ∼240 million years ago in the Mid-Triassic, to the crocodilian ancestor, which existed ∼80 million years ago in the Late Cretaceous (Fig. 1*A*).

**Figure 1.**
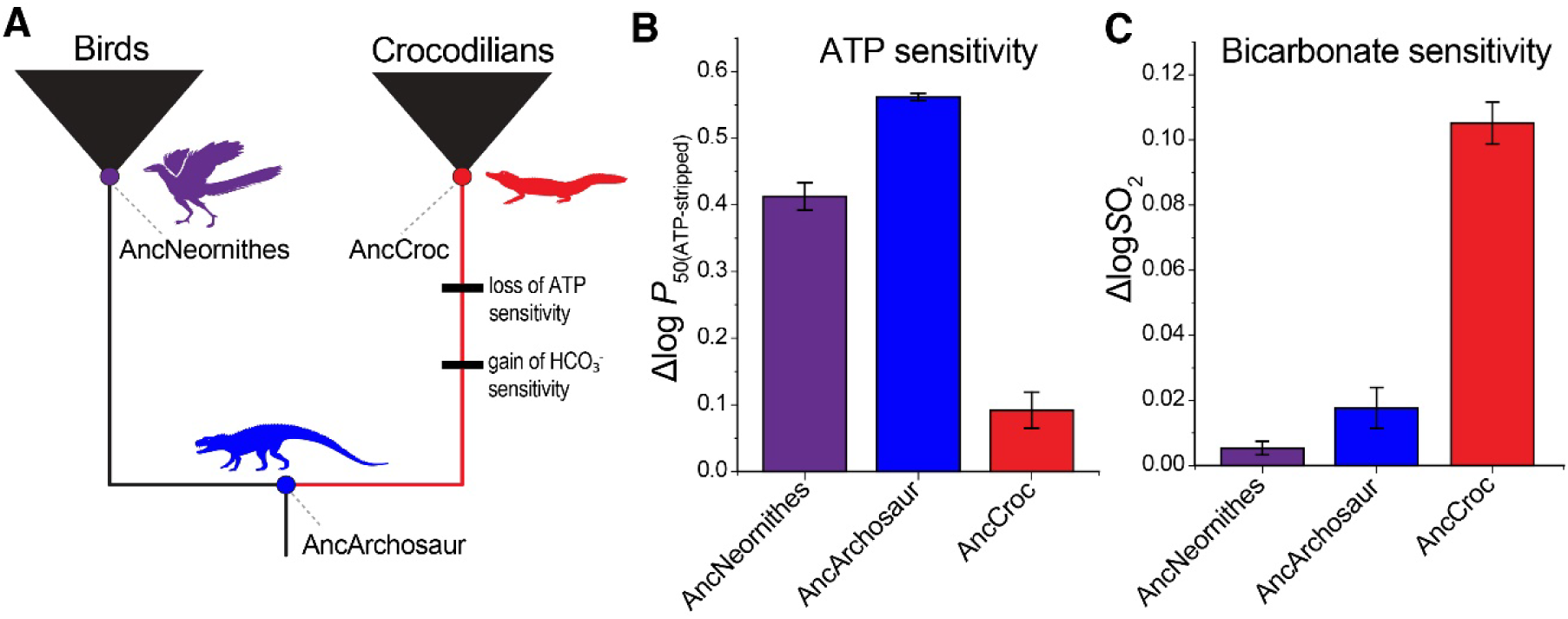
Resurrection and functional testing of ancestral hemoglobins. (***A***) Phylogeny of extant archosaurs, indicating nodes targeted for ancestral protein resurrection. The red branch connecting the common ancestor of modern archosaurs (AncArchosaur) with the ancestor of crocodilians (AncCroc) brackets the interval during which crocodilian Hb evolved the capacity to regulate Hb-O_2_ affinity via allosteric binding of bicarbonate ions. The gain of bicarbonate-sensitivity was accompanied by a loss of ATP-sensitivity (the ancestral mode of allosteric regulatory control in vertebrates). Relative to AncNeornithes and AncArchosaur, AncCroc evolved (***B***) a derived reduction in ATP-sensitivity (measured as the differences in log-transformed values of *P*_50_ [the partial pressure of O_2_ at which Hb is half-saturated] in presence and absence of 0.2 mM ATP [25° C, pH 7.4]) and (***C***) an increase in bicarbonate sensitivity (measured as the difference in log-transformed values of *S*O_2_ [Hb-O_2_ saturation] in the presence and absence of 1% CO_2_). See *SI Appendix* for details. Graphed values are means based on triplicate measurements (±SEM).

## Results

### Ancestral protein resurrection

We used a maximum likelihood (ML) approach to estimate the α- and β-globin sequences of three vertebrate ancestors: the most recent common ancestor of archosaurs (‘AncArchosaur’), the ancestor of modern birds (the sister group of crocodilians; ‘AncNeornithes’), and the ancestor of modern crocodilians (‘AncCroc’)(Fig. 1*A* and Figs. S1-S2). The ML ancestral sequences were estimated with high levels of statistical confidence (Fig. S3*A*-*C*).

We synthesized the ML α- and β-chain sequences of AncArchosaur, AncNeornithes, and AncCroc and cloned them into custom-designed expression plasmids (15, 16). We expressed and purified each of the three recombinant Hbs (rHbs), and we then measured oxygenation properties *in vitro*. The reconstructed AncArchosaur and AncCroc rHbs exhibited intrinsic O_2_-affinities, cooperativities, and pH sensitivities (Bohr effect) that are typical of native Hbs from modern crocodilians and non-avian reptiles (4, 6, 17-20)(Table S1). We used an experimental approach to confirm that all functional inferences are robust to uncertainty in ancestral sequence estimates (21)(*Supplementary Text* and Fig. S3*D*-*F*). The key finding is that AncArchosaur and AncNeornithes rHbs exhibited sensitivity to ATP but not to bicarbonate; conversely, the AncCroc rHb exhibited sensitivity to bicarbonate but not to ATP (Fig. 1*B*-*C*). These results confirm that AncArchosaur and AncCroc bracket an evolutionary transition between two discrete, well-defined functional states.

### Molecular changes underlying the loss of ATP sensitivity

Results of structural modeling suggest that ATP-binding to the T-state of vertebrate Hb is mediated by direct interactions with a constellation of positively charged residues in the central cavity (4, 22, 23). Structural comparisons between AncArchosaur and AncCroc Hbs suggested that the croc-specific loss of ATP-sensitivity may have been caused by substitutions at a total of five β-chain sites: 2, 82, 135, 139, and 143. We therefore introduced croc-specific amino acid states at each site on the AncArchosaur background, individually and in combination (‘AncArchosaur+5’). Four of the five mutations produced modest reductions in ATP sensitivity when introduced individually on the AncArchosaur background, but the net effect of all five mutations did not reduce ATP sensitivity to the same level as AncCroc Hb (Fig. 2*A*). We therefore introduced two additional croc-specific substitutions (Kβ87N and Kβ144E) that eliminated positive charges in the central cavity. While neither Kβ87 nor Kβ144 directly bind phosphate (4, 22), we hypothesized that the reduction in net positive charge caused by croc-specific substitutions at these two sites might further inhibit ATP-binding. Consistent with this prediction, adding these two charge-changing mutations to the ‘AncArchosaur+5’ construct (‘AncArchosaur+7’) reduced ATP sensitivity to the same level as AncCroc (Fig. 2*A*). Thus, a total of seven β-chain substitutions in the central cavity (including two that are not directly involved in phosphate-binding) are sufficient to explain the evolved loss of ATP-sensitivity in the ancestor of crocodilians. The substitutions that were sufficient to eliminate ATP-sensitivity did not simultaneously enhance bicarbonate sensitivity to the level observed in AncCroc (Fig. S4*A*), indicating that the gain and loss of different allosteric interactions involve different substitutions.

**Figure 2.**
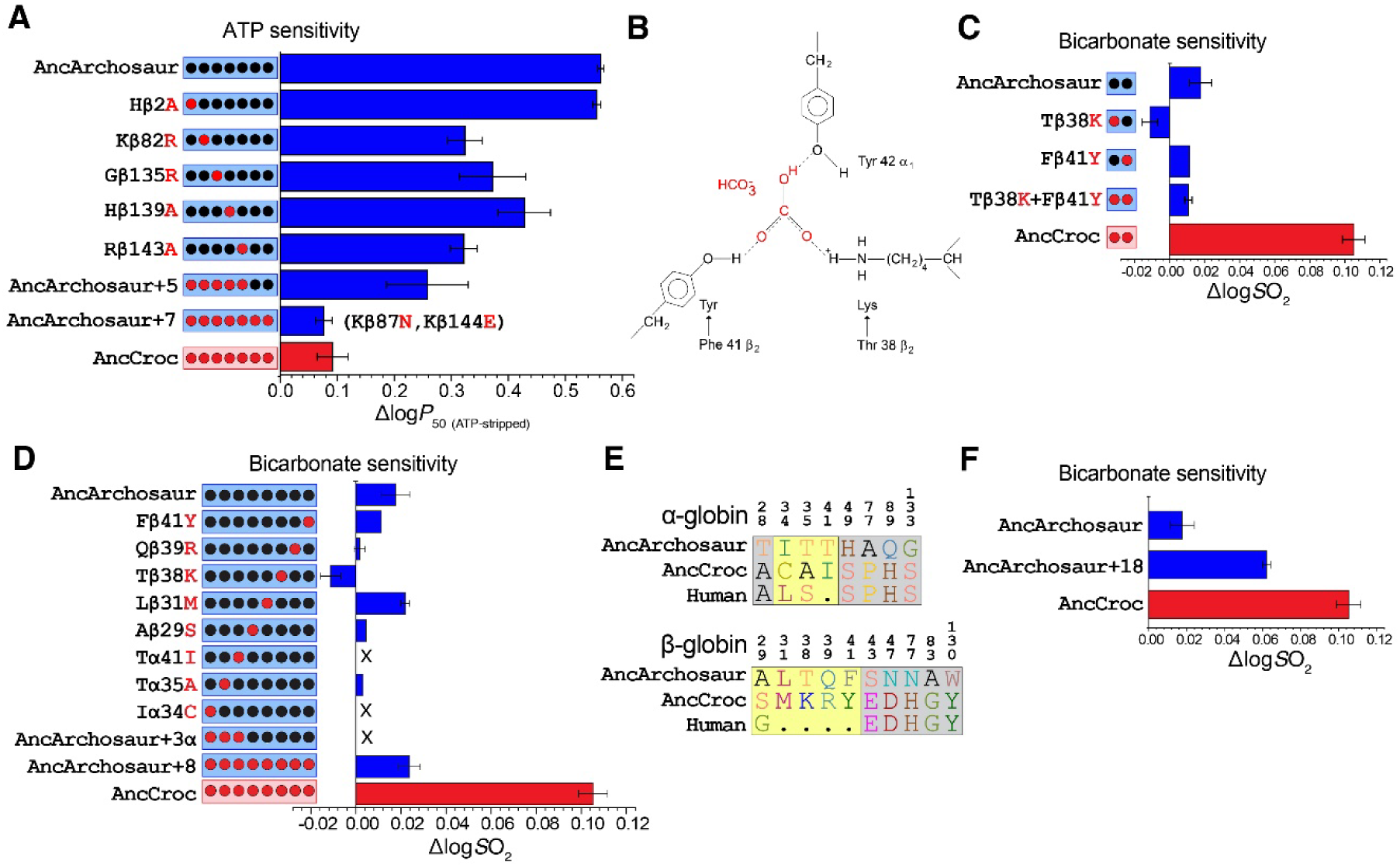
Experimental tests of hypotheses regarding the amino acid substitutions responsible for the loss of ATP-sensitivity and gain of bicarbonate-sensitivity in the ancestor of crocodilians. **(*A*)** A total of seven croc-specific substitutions is sufficient to eliminate ATP-sensitivity. Two of the substitutions, Kβ87N and Kβ144E, reduce the net positive charge in the central cavity but are not directly involved in phosphate-binding. **(*B*)** Perutz’s hypothesized binding site for bicarbonate ion (HCO_3_ ^-^) at the symmetrical α_1_β_2_ and α_2_β_1_ interdimeric interfaces of the Hb tetramer. Substitutions Tβ38K and Fβ41Y both occurred in the stem lineage of crocodilians, whereas Tyr-α42 is conserved among all modern archosaurs. **(*C*)** Contrary to the hypothesized mechanism, introducing both croc-specific mutations on the AncArchosaur background – individually and in combination –did not augment bicarbonate sensitivity. **(*D*)** Effects of individual mutations at 8 candidate sites on bicarbonate-sensitivity. When introduced individually on the AncArchosaur background, two α-chain mutations, I34C and T41I (located in α_1_β_1_/α_2_β_2_ and α_1_β_2_/α_2_β_1_ interfaces, respectively), resulted in nonfunctional protein (denoted by an ‘X’). The same was true when both mutations were combined with T35A as part of a trio of croc-specific α-chain mutations (‘AncArchosaur+3α’). None of the 8 candidate mutations significantly increased bicarbonate sensitivity above the AncArchosaur reference value, individually or in combination (‘AncArchosaur+8’). **(*E*)** The set of candidate mutations for bicarbonate sensitivity include eight that were previously tested by Komiyama *et al*. (11)(yellow background) in combination with 10 possible permissive substitutions where AncCroc and human share amino acid states to the exclusion of AncArchosaur (grey background). **(*F*)** The combination of 18 mutations on the AncArchosaur background produced a level of bicarbonate sensitivity equal to 59% of the AncCroc reference value. Graphed values are means based on triplicate measurements (±SEM).

### Molecular changes underlying the gain of bicarbonate sensitivity

In considering the biophysical mechanism of bicarbonate-binding to crocodilian Hb, Perutz (24) suggested that two derived (croc-specific) amino acids, Kβ38 and Yβ41, bind a single bicarbonate ion in coordination with the highly conserved tyrosine at α42 (located on the α-chain subunit of the opposing αβ dimer)(Fig. 2*B*). This mechanism predicts the binding of two bicarbonate ions per Hb tetramer, one at each of the two symmetrical interdimer interfaces (α_1_β_2_ and α_2_β_1_), and is consistent with experimental results for native crocodilian Hbs which documented that bicarbonate ions bind deoxyHb with a 2:1 stoichiometry (1, 6). To test the Perutz hypothesis, we synthesized four rHbs representing the requisite single- and double-mutant genotypes and measured the individual and combined effects of croc-specific mutations (Tβ38K and Fβ41Y) on the AncArchosaur background. Contrary to Perutz’s prediction, introduction of croc-specific mutations at sites β38 and β41– individually and in pairwise combination – did not increase bicarbonate sensitivity relative to the ancestral reference value (Fig. 2*C*). We then expanded the focus to the set of 12 amino acid states that Komiyama *et al*. (11) had identified as being necessary and sufficient to transplant bicarbonate sensitivity into human Hb, but we engineered these same changes into the AncArchosaur background. Since AncArchosaur and AncCroc share the same amino acid state at four of the 12 sites mutagenized by Komiyama *et al*., we only needed to introduce a total of eight new mutations on the AncArchosaur background. Results revealed that the eight mutations – individually and in combination – did not significantly increase bicarbonate sensitivity (Fig. 2*D*), nor did they significantly alter ATP-sensitivity (Fig. S4*B*).

Why were the set of 12 croc-specific mutations sufficient to confer the bicarbonate effect in human Hb, but not in AncArchosaur Hb? One possible explanation is that one or more substitutions specific to human Hb may potentiate bicarbonate sensitivity in combination with croc-specific amino acid states at the 12 focal sites. To nominate candidates for such permissive substitutions, we identified 10 sites where human and AncCroc share the same amino acid state to the exclusion of AncArchosaur. We then synthesized a new version of AncArchosaur (‘AncArchosaur+18’) that contained the eight croc-specific substitutions tested by Komiyama *et al*. in combination with the 10 candidate permissive substitutions (Fig. 2*E*). This set of 18 changes produced a significant increase in bicarbonate sensitivity, but it was still only 59% of the reference value for AncCroc Hb (Fig. 2*F*). We then investigated an alternative biophysical mechanism by testing different multi-site combinations of 19 croc-specific substitutions that alter the net charge of the central cavity and/or alter proton dissociation constants of Lys or Arg sidechains that could serve as binding sites for bicarbonate ions at a close-to-neutral pH (*Supplementary Text*). None of the tested combinations recapitulated the full bicarbonate-sensitivity of AncCroc (Fig. S5).

### Insights from Hb isoform differentiation

The evolved transition between the bicarbonate-insensitive ancestor (AncArchosaur) and the bicarbonate-sensitive descendant (AncCroc) is the rationale for testing croc-specific forward mutations on the ancestral AncArchosaur background. Since none of our mutagenesis experiments on the AncArchosaur background successfully recapitulated the full bicarbonate sensitivity of AncCroc, we explored an alternative approach that enabled us to isolate the functional effect of β-chain substitutions. Crocodilians possess multiple copies of β-type globin genes (25) and express structurally distinct Hb isoforms (isoHbs) at different stages of prenatal development, as is the case with all other amniotes (22, 26-28). We previously discovered that crocodilian embryos express two structurally distinct isoHbs: an embryonic isoHb that incorporates the β-chain product of *HBB-T1* (HbI) and the normal adult isoHb that incorporates the β-chain product of *HBB-T4* (HbII); both isoHbs share the same α-chain subunit (20)(Fig. 3*A*). *HBB-T1* and *HBB-T4* are products of a croc-specific duplication event (4, 25), so it is possible that embryonic HbI (with a β-chain subunit distinct from that of the adult HbII) retains the ancestral bicarbonate-insensitive condition typical of all other amniote Hbs. If so, examination of isoform differences between embryonic HbI and adult HbII could provide a tractable means of identifying the molecular basis of bicarbonate sensitivity. To investigate isoHb differences in bicarbonate sensitivity, we isolated and purified HbI and HbII in the red blood cells of alligator embryos (*Alligator mississippiensis*) at day 40 of pre-hatching development. We experimentally confirmed that adult HbII is sensitive to bicarbonate but not to ATP (Fig. 3*B*), as expected based on previous experiments on native Hbs purified from red blood cells of adult crocodilians (4, 6). Importantly, we discovered that the embryonic HbI is *not* sensitive to bicarbonate, but *is* sensitive to ATP, just like AncArchosaur and the native Hbs of birds and all other amniotes (Fig. 3*B*). Since the embryonic and adult isoHbs share the same α-chain, the observed isoHb differences indicate that bicarbonate-sensitivity can evolve from an insensitive ancestral state via β-chain substitutions alone.

**Figure 3.**
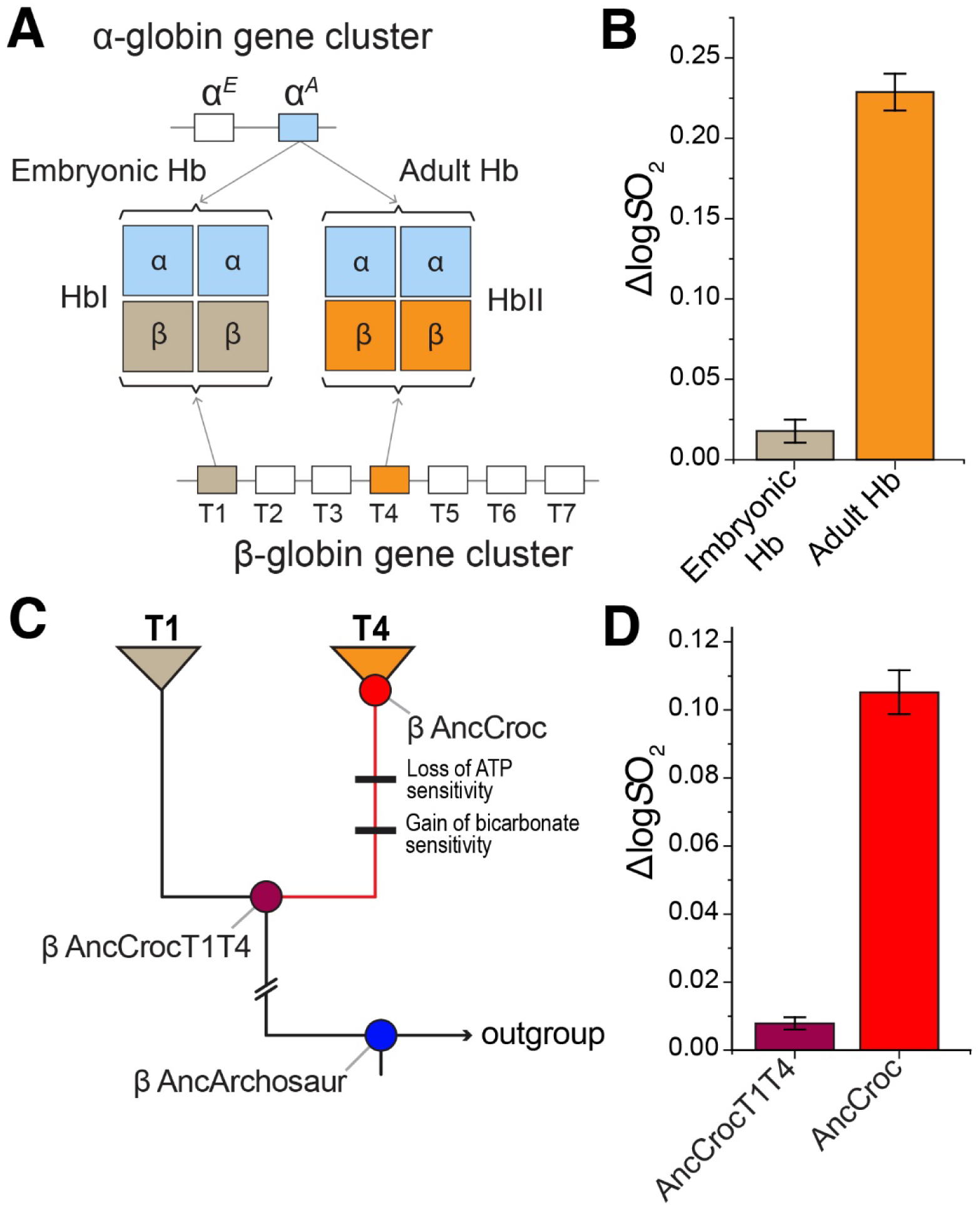
Evolved differences in allosteric properties of embryonic and adult crocodilian Hb isoforms provide an alternative means of investigating the molecular basis of bicarbonate sensitivity. **(*A*)** Embryonic and adult isoHbs of crocodilians: HbI and HbII, respectively. The two isoHbs share identical α-chains but incorporate structurally distinct β-chain products of the *HBB-T1* and *HBB-T4* paralogs. **(*B*)** In contrast to the adult HbII isoform, embryonic HbI is insensitive to bicarbonate (the ancestral condition, shared by Hbs of all other amniotes). **(*C*)** Phylogeny of croc-specific β-globin paralogs showing internal nodes targeted for ancestral protein resurrection. **(*D*)** Experiments on resurrected ancestral proteins confirmed that the β-chain substitutions that conferred bicarbonate sensitivity occurred in the post-duplication branch (red) that connects ‘β AncCrocT1/T4’ (the single-copy, pre-duplication ancestor of the croc-specific *HBB-T1* and *HBB-T4* paralogs) to ‘β AncCroc’. β AncCrocT1/T4 and AncCroc therefore bracket an evolutionary transition between ancestral and derived functional states. Graphed values are means based on triplicate measurements (±SEM).

To winnow down the list of candidate β-globin mutations for bicarbonate sensitivity, we estimated the pre-duplication ancestor of the croc-specific *HBB-T1* and *HBB-T4* paralogs (β AncCrocT1/T4) (Fig. 3*C* and Fig. S6), which enabled us to conduct a vertical comparison between the bicarbonate-insensitive Hb that incorporates the single-copy progenitor of *HBB-T1*/*T4* and the bicarbonate-sensitive Hb that incorporates the product of *HBB-T4* (=β AncCroc)(Fig. 3*C*-*D*). As in the case with the AncArchosaur vs. AncCroc comparison, AncCrocT1/T4 vs. AncCroc also constitutes a vertical comparison between a bicarbonate-insensitive ancestor and a bicarbonate-sensitive descendant, but the ancestral and descendant genotypes differ at fewer sites (49 vs. 74) and all substitutions are restricted to the β-chain. As a first step, we introduced a total of 13 AncCroc-specific mutations into the β AncCrocT1/T4 background: the eight β-chain mutations tested by Komiyama *et al*. in combination with five charge- or polarity-changing mutations concentrated in the α_1_β_2_/α_2_β_1_ interface (Fig. S7), including the Tβ38K and Fβ41Y mutations that are central to the Perutz hypothesis (Fig. 2*B*). This combination of mutations (‘AncCrocT1/T4+13’) produced a significant increase in bicarbonate-sensitivity, reaching 61% of the AncCroc reference value (Fig. 4*A*). Reverting these mutations to the ancestral amino acid states on the AncCroc background eliminated bicarbonate sensitivity (Fig. 4*A*). Follow-up experiments revealed that the increased bicarbonate sensitivity could be explained by the five croc-specific α_1_β_2_/α_2_β_1_ interface mutations alone (‘AncCrocT1/T4+5’, which included Tβ38K and Fβ41Y) and, again, reverting those same five mutations on the AncCroc background was sufficient to eliminate sensitivity (Fig. 4*A*). Although the ‘+5’ set of mutations in the α_1_β_2_/α_2_β_1_ interface may be necessary to confer bicarbonate sensitivity, those five changes alone are clearly not sufficient to recapitulate the full sensitivity of AncCroc. To test whether some combination of other croc-specific β-chain mutations need to be combined with the +5 mutations to confer full sensitivity, we divided the AncCrocT1/T4 β-chain sequence into thirds, and we synthesized rHb constructs containing all croc-specific substitutions in the first, second, and third parts (residues 1-49, 50-98, and 99-146, respectively). The ‘Part I’ construct contained the ‘+5’ mutations in combination with 16 additional croc-specific mutations concentrated in the N-terminus and α_1_β_2_/α_2_β_1_ and α_1_β_1_/α_2_β_2_ interfaces. Experimental results demonstrate that the set of 21 croc-specific ‘Part I’ mutations successfully recapitulates the full bicarbonate sensitivity of AncCroc (Fig. 4*B*).

**Figure 4.**
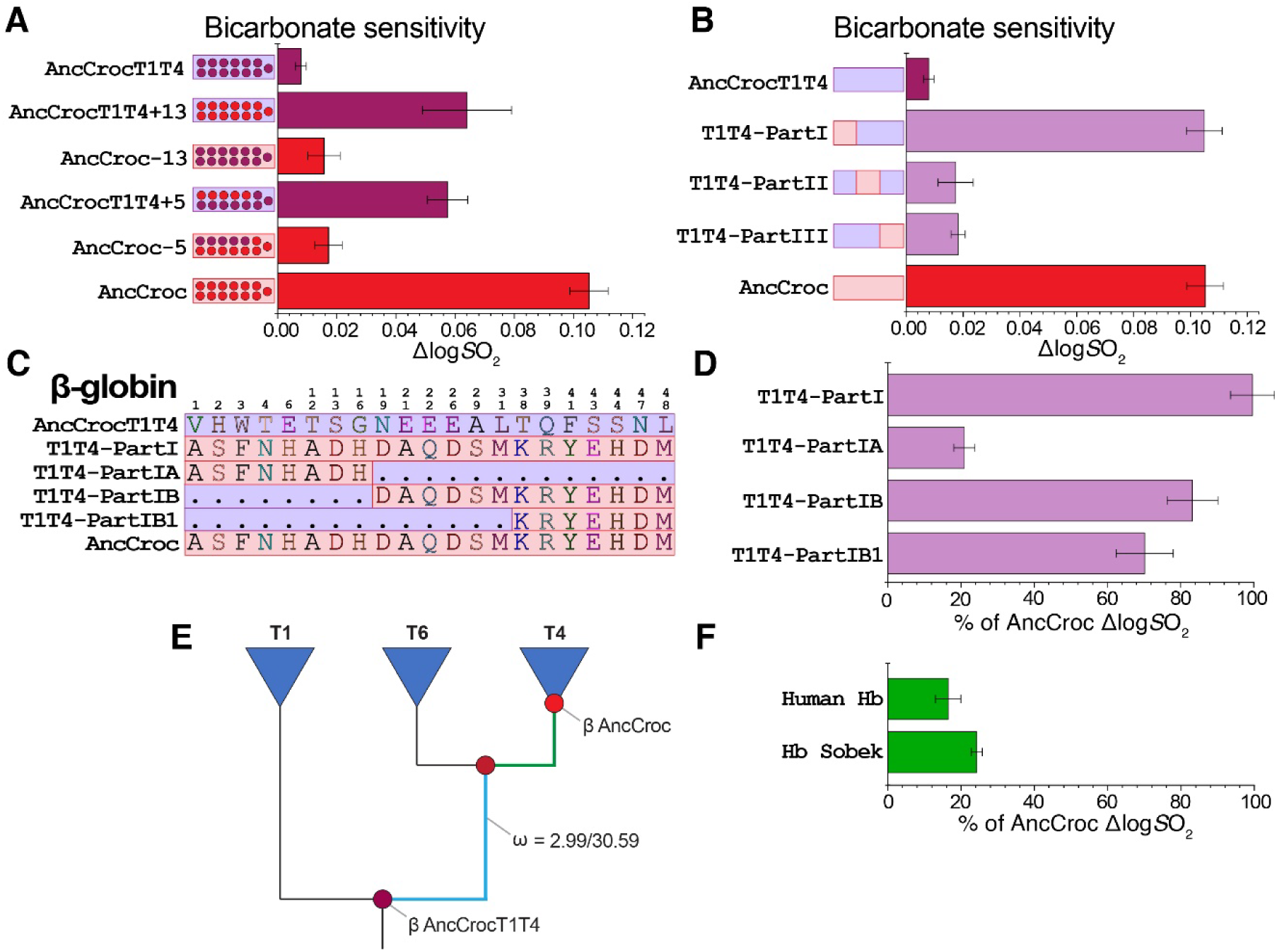
Testing candidate mutations for bicarbonate sensitivity on the AncCrocT1/T4 background. **(*A*)** At 13 candidate sites, forward mutations on the AncCrocT1/T4 background (blue) produce a significant enhancement of bicarbonate-sensitivity that falls short of the AncCroc reference value. The observed enhancement can be explained by the effects of just five croc-specific mutations in the α_1_β_2_/α_2_β_1_ interface. Reverting such mutations on the AncCroc background (red) eliminates bicarbonate sensitivity. **(*B*)** To test whether some combination of other croc-specific β-chain mutations need to be combined with the +5 mutations to confer full sensitivity, we divided the AncCrocT1/T4 β-chain sequence into thirds, and we tested constructs containing all croc-specific substitutions in each of the three parts. ‘Part 1’ contained the ‘+5’ mutations in combination with 16 additional croc-specific mutations concentrated in the N-terminus and α_1_β_2_/α_2_β_1_ and α_1_β_1_/α_2_β_2_ interfaces. Results demonstrate that the set of 21 croc-specific ‘Part I’ mutations successfully recapitulates the full bicarbonate sensitivity of AncCroc. **(*C*)** We synthesized additional constructs representing further subdivisions of ‘Part I’. **(*D*)** The smaller subsets of Part I mutations do not fully recapitulate bicarbonate sensitivity, although the set of mutations in the α_1_β_2_/α_2_β_1_ interface (including Tβ38K and Fβ41Y) make the largest contribution. **(*E*)** Diagrammatic phylogeny of crocodilian β-globin paralogs showing a significantly accelerated rate of nonsynonymous substitution (ω [=*d*_N_/*d*_S_] in branch model/clade model) in the post-duplication branch connecting the single-copy progenitor of the *HBB-T1* and *HBB-T4* paralogs (β AncCrocT1/T4, which is also the common ancestor of all crocodilian β-globins) to the most recent common ancestor of *HBB-T4* orthologs from all extant crocodilians (β AncCroc). **(*F*)** No significant difference in bicarbonate sensitivity between human Hb and the mutant Hb Sobek that contains the same 21 croc-specific substitutions that were sufficient to confer full bicarbonate sensitivity on the AncCrocT1/T4 background (T1T4-PartI). Graphed values in panels *A, B, D*, and *F* are means based on triplicate measurements (±SEM)

Having identified a set of 21 croc-specific substitutions that are sufficient to confer full bicarbonate sensitivity on the AncCrocT1/T4 background, we then performed additional mutagenesis experiments in an attempt to further winnow the number of necessary substitutions. We synthesized new β AncCrocT1/T4 constructs that divided Part I into two equal halves, Part IA and Part IB (containing all croc-specific mutations between β-chain residues 1-18 [positions NA1 to AB1] and 19-49 [positions C1 to CD8], respectively), and PartIB1 (which contains a subset of 7 croc-specific substitutions in Part IB that span the α_1_β_2_/α_2_β_1_ interface)(Fig. 4*C*). Experimental results revealed that the set of croc-specific substitutions concentrated at the β-chain N-terminus (Part IA) produced no significant effect, whereas the 13 croc-specific mutations in Part IB conferred a level of bicarbonate sensitivity equal to 83.3 % of the reference value for AncCroc Hb (Fig. 4*D*). When that set of 13 croc-specific mutations was pared down to a subset of 7 mutations in α_1_β_2_/α_2_β_1_ interface (Part IB1), bicarbonate-sensitivity decreased to 70.2 % of the reference value (Fig. 4*D*). It is clear that full bicarbonate sensitivity requires numerous substitutions that are distributed across multiple subdomains of the protein.

In summary, mutagenesis experiments revealed a set of 21 β-chain substitutions that are sufficient to confer full bicarbonate sensitivity on the ancestral AncCrocT1/T4 background, and a small subset of substitutions in the α_1_β_2_/α_2_β_1_ interface make a major contribution. In contrast to results of previous efforts to transplant bicarbonate-sensitivity into human Hb (11), introducing our set of 21 croc-specific mutations onto the AncCrocT1/T4 background successfully conferred bicarbonate-sensitivity without compromising intrinsic O_2_-affinity or cooperativity (Table S1). The two croc-specific substitutions highlighted by Perutz, Tβ38K and Fβ41Y, appear to be necessary to confer the bicarbonate effect on the T1/T4 background, but those two substitutions alone are not sufficient. The full sensitivity observed in AncCroc is only achieved when the interface mutations are combined with numerous other croc-specific amino acid states within the N-terminal 1/3 of the β-chain. It may be that the triad of residues Kβ38, Yβ41, and Tα42 constitute the only site for oxygenation-linked binding of bicarbonate (Fig. 2*B*), but the requisite stereochemical conformation of the three residues may depend on a higher-level (quaternary) conformation that is determined by the amino acid states of structurally remote sites.

### Tests of positive selection

Given that the gain of bicarbonate sensitivity in crocodilian Hb is attributable to β-chain substitutions that occurred in the adult β-globin gene (*HBB-T4*) following duplication of the single-copy ancestor of *HBB-T1* and *HBB-T4* (β AncCroc T1/T4), we performed a molecular evolution analysis to test for evidence of positive selection in the post-duplication branch of the gene tree that connects β AncT1/T4 to β AncCroc (Fig. S6). This test revealed evidence for a strikingly accelerated rate of nonsynonymous substitution in the branch that connects β AncT1/T4 to the single-copy, pre-duplication ancestor of the croc-specific paralogs *HBB-T4* (= β AncCroc) and *HBB-T6* (which may encode β-chain subunits of an isoHb expressed in early embryogenesis) (Fig. 4E and Table S2), and a clade test revealed significantly accelerated substitution rates among *HBB-T4* orthologs of different crocodilian lineages (Table S3). This molecular evolution analysis provides an independent line of evidence that the bicarbonate-sensitivity of crocodilian Hb represents an adaptive trait that evolved via strong positive selection.

### Can bicarbonate-sensitivity be transplanted into human Hb

Having discovered a set of 21 croc-specific substitutions that are sufficient to confer bicarbonate-sensitivity when introduced into ancestral crocodilian Hb, we tested whether the same set of replacements produce the same phenotypic effect when introduced into a divergent genetic background. Specifically, we used site-directed mutagenesis to introduce the 21 ‘Part I’ croc-specific mutations into recombinant human Hb (‘Hb Sobek’, named after the half-crocodile/half-human Egyptian deity). The experiment revealed that the 21 mutations that were sufficient to confer bicarbonate-sensitivity on the crocodilian genetic background produced a negligible increase in sensitivity when introduced into the structural context of human Hb and did not come close to the full allosteric regulatory capacity of AncCroc (Fig. 4*F*).

## Discussion

Our experimental results provide answers to several fundamental questions about protein evolution and biochemical adaptation. First: Are adaptive changes in protein function attributable to few substitutions at key sites, or many substitutions that have individually minor effects? Perutz (3) suggested that adaptive modifications of Hb function will typically be attributable to ‘…a few replacements in key positions’, a scenario consistent with the evolution of substrate-specificity in metabolic enzymes (29) and ligand-specificity of allosterically regulated steroid receptors (30, 31). With regard to the loss of ATP-sensitivity, we identified several croc-specific mutations that directly eliminated phosphate-binding sites, but the full loss of sensitivity required the indirect effect of two additional mutations that altered the net positive charge in the central cavity. With regard to the gain of bicarbonate-sensitivity, we identified two croc-specific mutations in the α_1_β_2_/α_2_β_1_ interface (Tβ38K and Fβ41Y) that appear to play a direct role in the allosteric binding of bicarbonate ion, but they confer their allosteric effect only in conjunction with mutations at numerous other sites that are structurally remote from the intersubunit interface. Thus, evolved changes in allosteric regulation involved the direct effect of few substitutions at key positions in combination with indirect effects of numerous other substitutions at structurally disparate sites. Such indirect interaction effects suggest that the evolution of novel functions may be highly dependent on neutral mutations that produce no adaptive benefit when they first arise but which contribute to a permissive background for subsequent function-altering mutations at other sites.

A second key question concerns the extent to which ancestral protein functions can be maintained during the evolutionary transition to a novel function. With regard to the evolution of crocodilian Hb, the question is whether it is possible to maintain the capacity for allosteric regulatory control by ATP-binding (the ancestral condition for the Hb of jawed vertebrates) while simultaneously evolving bicarbonate sensitivity. Results of our experiments indicate that it is possible to maintain sensitivity to both ATP and bicarbonate, and that there is no ineluctable trade-off between the two modes of allosteric regulation. The minimal set of amino acid mutations that are sufficient to eliminate ATP sensitivity on the AncArchosaur background do not simultaneously affect bicarbonate sensitivity. Likewise, none of the mutations implicated in the gain of bicarbonate sensitivity affect allosteric ATP-binding.

A third key question concerns the context-dependence of mutations (epistasis) and its implications for adaptive protein evolution. Our experimental attempt to transplant bicarbonate-sensitivity into human Hb provides a striking example of intramolecular epistasis: the same 21 mutations that conferred full bicarbonate sensitivity when introduced into ancestral crocodilian Hb produced no effect in human Hb. This context-dependence highlights the importance of using a ‘vertical’ approach in mutagenesis experiments (introducing forward mutations on ancestral backgrounds) to identify evolved mechanisms of functional change (12). The result also suggests an important role for historical contingency in the evolution of novel protein functions, as it indicates that some adaptive solutions are not equally accessible from all possible ancestral starting points. The fact that causative substitutions in crocodilian Hb have no effect on the divergent genetic background of human Hb suggest that bicarbonate-sensitivity may not be a mutationally accessible design solution for Hbs of other vertebrates, or that the same allosteric property would have to evolve via a completely different molecular mechanism.

## Materials and Methods

### Estimation of ancestral globin sequences

We estimated α- and β-type globin sequences of ancestral Hbs representing three internal nodes in the archosaur phylogeny (AncNeornithes, AncArchosaur, and AncCroc) (*Supplementary Material*, Figs. S1 and S2). For each analysis, we collected a set of non-redundant α- and β-globin sequences that provided balanced phylogenetic coverage of extant archosaur diversity. We estimated all ancestral sequences using the maximum likelihood (ML) approach implemented in PAML version 4.8 (32), with the aid of the Lazarus set of Python scripts for parsing results (33). See *Supplementary Material* for details. All reconstructed sequences that we used for ancestral protein resurrection were deposited in GenBank (ON959522-ON959565).

### Vector construction and site-directed mutagenesis

The reconstructed amino acid sequences of AncArchosaur, AncNeornithes, AncCroc, and AncCrocT1T4 were reverse-translated to DNA sequence and were optimized for *E. coli* codon preferences. Gene cassettes were subcloned into the pGM custom expression system as described previously (15, 34-36). See *SI Appendix* for details.

### Protein expression and purification

Expression of recombinant Hbs (rHbs) was carried out in the JM109(DE3) *E. coli* strain (New England Biolabs). To ensure that N-terminal methionines were post-translationally cleaved from the nascent globin chains, host cells were co-transformed with a plasmid (pCOMAP) that contained an additional copy of the methionine aminopeptidase gene (MAP) along with a kanamycin resistance gene (15, 16). See *Supplementary Material* for details.

### Collection of blood samples and purification of native isoHbs

We sampled blood of alligator embryos at the 40% stage of pre-hatching development by catheterizing the major artery of the chorioallantoic membrane, as described by Bautista *et al*. (20). See *Supplementary Material* for details regarding hemolysate preparation and protein purification via anion-exchange chromatography.

### Measurement of oxygen equilibration curves

Solutions of purified rHbs were desalted by passing samples through PD-10 desalting columns (GE Healthcare) that were equilibrated with 0.01 M HEPES, 0.5 mM EDTA, pH 7.4, and concentrated using Amicon Ultra-4 Centrifugal Filter Units (MilliporeSigma). O_2_-equilibrium curves for Hb solutions (0.4 mM Hb tetramer in 100 mM HEPES, 0.5 mM EDTA buffer) were measured at 25**°**C using a Blood Oxygen Binding System (BOBS; Loligo^®^ Systems). Each Hb solution was sequentially equilibrated with 3-5 different partial pressures of O_2_ (*P*O_2_’s) at saturation levels between 30 to 70%, while absorbance was continually monitored at 430 nm (deoxy peak) and 421 nm (oxy/deoxy isosbestic point). Hb-O_2_ saturation was estimated at each equilibrium step by comparing the absorbance at 430 nm to fully oxygenated and deoxygenated baselines. Estimates of the *P*O_2_ at half-saturation (*P*_50_) and the cooperativity coefficient (*n*_50_) were then estimated from linear Hill plots (log[fractional saturation/[1-fractional saturation]] vs. log*P*O_2_). O_2_-equilibrium curves were measured in the absence (stripped) and presence of Cl^-^ ions (0.1 M KCl) and organic phosphates (0.2 mM adenosine triphosphate [ATP]). O_2_-equilibrium curves were also measured at three different pH levels, where the pH of working solutions was adjusted with NaOH to 7.2, 7.4, or 7.6. Bicarbonate sensitivity of the ancestral and mutant rHbs was measured using 3 µL samples (0.14 mM heme, 0.1 M HEPES, pH 7.2, 0.5 mM EDTA at 25°C)(38-40). See *Supplementary Material* for details.

### Tests of positive selection

Using a maximum likelihood (ML) framework, we tested for evidence of positive selection in the crocodilian β-globin genes with codon-based models, as implemented in the *codeml* program in PAML v4.9 (38). We estimated phylogenetic relationships among crocodilian β-globin genes using IQTree version 2.1.3 (39) under the best-fitting model selected by the ModelFinder subroutine in IQ-Tree (40) and we used chicken β^*A*^-globin as an outgroup (Fig. S6). We used the branch-site and clade models to examine variation in ω, the ratio of the rate of nonsynonymous substitution per nonsynonymous site, *d*_N_, to the rate of synonymous substitution per synonymous site, *d*_S_. Using both the branch and branch-site models (41, 42), we tested for changes in ω in the post-duplication line of descent leading from the single-copy progenitor of the *HBB-T1* and *HBB-T4* genes (β AncCrocT1/T4, which is also the common ancestor of all crocodilian β-type globin paralogs) to the most recent common ancestor of *HBB-T4* orthologs from all extant crocodilians (= the β-chain of AncCroc Hb)(Fig. S6). See *Supplementary Material* for details.

## Acknowledgments

We thank A. Kumar and E. Petersen for assistance in the lab. This work was funded by National Institutes of Health (NIH) grant R01HL087216 (J.F.S.) and National Science Foundation (NSF) grant OIA-1736249 (J.F.S.).

