## Supplementary Material for "Evolution and molecular basis of a novel allosteric property of crocodilian hemoglobin"

**This PDF file includes:**

Supporting Information Text
Figures S1 to S7
Tables S1 to S4
References

### Supporting Information Text

#### Results

**Testing whether inferences about evolved functional properties are robust to statistical uncertainty in ancestral sequence estimates**. To assess the robustness of our functional inferences to uncertainties in ancestral sequence estimates, we synthesized alternative versions of the AncArchosaur, AncNeornithes, AncCroc, and AncCrocT1/T4 recombinant Hbs (rHbs) that incorporated the alternative amino acid at each ambiguous site in the corresponding ML sequences (1)(Fig. S3*A*-*C*). Specifically, we synthesized 'AltAll' sequences by incorporating the 'runner-up' amino acid at each site with a posterior probability <0.80. In the case of AncArchosaur, the AltAll sequences differed from the ML ancestral sequences at 19 sites (18 in the α-chain and 1 in the β-chain). In the case of AncNeornithes and AncCroc, AltAll sequences differ from the corresponding ML sequences at 4 α-chain sites and 11 β-chain sites, respectively. In the case of β AncCrocT1/T4, the AltAll sequence differed from the corresponding ML sequence at 5 sites. With regard to ancestral sequence estimates, the AltAll sequences can be interpreted as worst case scenarios (1). If the ML ancestor and the corresponding AltAll protein have similar functional properties, this provides evidence that the inferred properties of the proteins are robust to statistical uncertainties in the ancestral sequence reconstruction.

The comparison between AncArchosaur and AncArchosaurAltAll revealed that the two proteins have highly similar oxygenation properties, and – in contrast to AncCroc – they both exhibit the expected sensitivity to ATP and neither are sensitive to bicarbonate (Fig. S3*D*, Table S1). The same was true for the comparisons AncNeornithes vs. AncNeornithesAltAll and AncCrocT1/T4 vs. AncCrocT1/T4AltAll (Fig. S3*E*, Table S1). Likewise, the comparison between AncCroc and AncCrocAltAll confirmed that these two proteins share similar allosteric properties (low ATP-sensitivity, high bicarbonate-sensitivity)(Fig. S3F and Table S1) that distinguish them from AncArchosaur, AncNeornithes, and AncCrocT1/T4 (Fig. 1*B*-*C*, Fig. 3*D*, and Table S1). These results demonstrate that our inferences regarding evolved changes in allosteric properties are robust to statistical uncertainty in the ancestral sequence estimates.

**Mutagenesis experiments that altered net charge and/or proton dissociation constants**. Since mutagenesis experiments involving the initial set of candidate substitutions on the AncArchosaur background failed to recapitulate the bicarbonate-sensitivity of AncCroc (Fig. 2*C*-*F*), we designed another set of experiments to test an alternative biophysical mechanism. Homology modeling of AncCroc Hb revealed a higher net positive charge in the central cavity relative to AncArchosaur Hb (2). We hypothesized that some or all of the croc-specific substitutions that increase the positive net charge in the central cavity are necessary to decrease the proton dissociation constant (p$K_a$) of Lys or Arg sidechains, thereby permitting such residues to bind $CO_2$ or bicarbonate at a close-to-neutral pH (the amino group must be unprotonated to form carbamino). We tested this hypothesis by measuring the effects of charge-changing mutations at 9 sites that comprise three spatially discrete clusters in the central cavity or interdimeric ($\alpha_1\beta_2/\alpha_2\beta_1$) interface. We tested the net effect of all 9 croc-specific mutations on the AncArchosaur background ('AncArchosaur+9') and we also tested each cluster of charged mutations individually. Of the three charge clusters, only the trio of mutations comprising 'cluster 3' produced a significant increase in bicarbonate sensitivity (32% of the AncCroc reference value), but this effect disappeared when all nine charge-changing mutations were tested in combination (Fig. S5*A*). We then introduced the 'cluster 3' mutations in combination with 10 additional croc-specific mutations (seven of which are charge-changing) at structurally proximal residues in the $\alpha_1\beta_2/\alpha_2\beta_1$ intersubunit interface (Fig. S5*B*), but this 'AncArchosaur+13' combination of changes had no effect on bicarbonate sensitivity (Fig. S5*C*).

### Materials and Methods

**Estimation of ancestral globin sequences**. We estimated α- and β-type globin sequences of ancestral hemoglobins (Hbs) representing three internal nodes in the archosaur phylogeny (AncNeornithes, AncArchosaur, and AncCroc) (Figs. S1 and S2). We estimated all ancestral sequences using the maximum likelihood (ML) approach implemented in PAML version 4.8 (3), with the aid of the Lazarus set of Python scripts for parsing results (4). For each analysis, we collected a set of non-redundant α- and β-globin sequences that provided balanced phylogenetic coverage of extant archosaur diversity (Figs. S1 and S2). We constructed supertrees for avian sequences by starting with a backbone provided by the total-evidence phylogeny from Jarvis *et al.* (5). Sequences from all species could be unambiguously assigned to their appropriate branches in this backbone tree. Subtrees for avian sequences were taken from Jetz *et al*. (6), which was constructed using the Hackett *et al*. (7) backbone. Relationships among the major groups of sauropsids were based on the phylogeny in Green *et al.* (8) and assignments of orthology were based on Hoffmann *et al.* (9). In all cases, we used fully annotated globin genes from high-coverage genome assemblies along with newly generated sequences from a phylogenetically diverse set of crocodilian species (2).

To estimate the ancestral α-globin sequences of AncArchosaur and AncNeornithes Hbs, we included orthologous sequences from representatives of each of the major crocodilian and avian lineages (Fig. S1*A*) along with two testudine sequences as outgroups. To estimate the ancestral β-globin sequences of AncArchosaur and AncNeornithes Hbs, we expanded the phylogenetic coverage of paralogous β-type globins due to the history of lineage-specific duplication in both birds and crocodilians (9, 10). The avian β-globin paralogs include σ-, ε-, $\beta^{H}$-, and $\beta^{A}$-globin (10), and those of crocodilians include *HBB-T1*, *HBB-T4*, *HBB-T6*, and *HBB-T7* (2, 9). We included β-globin paralogs from *Anolis* and Chinese softshell turtle as outgroups (Fig. S1*B*). To estimate the ancestral α-globin sequence of AncCroc Hb, we expanded phylogenetic coverage of the adult-expressed crocodilian α-globin gene that is orthologous to avian $\alpha^{A}$-globin (9, 11, 12). To estimate the ancestral β-globin sequence of AncCroc Hb, we used an expanded supertree that included the full complement of croc-specific β-globin paralogs from representatives of each of the three extant families of crocodilians (Alligatoridae [alligators and caimans], Crocodylidae [true crocodiles], and Gavialidae [gharial and false gharial]). As outgroups, we included paralogous β-globin sequences from human, *Coelacanth*, *Xenopus,* and spotted gar (Fig. S2*B*). We used this same set of β-globin sequences to estimate the pre-duplication, single-copy ancestor of the crocodilian *HBB-T1* and *HBB-T4* paralogs ('β AncCrocT1/T4') (Fig. S6). All reconstructed sequences that we used for ancestral protein resurrection were deposited in GenBank (ON959522-ON959565).

**Vector construction and site-directed mutagenesis**. The reconstructed amino acid sequences of AncArchosaur, AncNeornithes, AncCroc, and AncCrocT1T4 were reverse-translated to DNA sequence and were optimized for *E. coli* codon preferences. The α- and β-globin gene cassettes were synthesized by GeneArt Gene Synthesis (Thermo Fisher Scientific). The gene cassettes were subcloned into the pGM custom expression system as described previously (13-18). Codon changes were engineered using the QuikChange II XL Site-Directed Mutagenesis kit (Agilent Technologies), with all mutated sites verified by Sanger sequencing.

**Protein expression and purification**. Expression of recombinant Hbs (rHbs) was carried out in the JM109(DE3) *E. coli* strain (New England Biolabs). To ensure that N-terminal methionines were post-translationally cleaved from the nascent globin chains, host cells were co-transformed with a plasmid (pCOMAP) that contained an additional copy of the methionine aminopeptidase gene (MAP) along with a kanamycin resistance gene (13, 17). Both pGM and pCOMAP plasmids were co-transformed and subject to dual antibiotic selection in an LB agar plate containing ampicillin and kanamycin. The expression of each rHb mutant was carried out in 1.5 to 2 L of TB medium. The cells were grown in an orbital shaker (New Brunswick™) at 37ºC, 200 rpm, until their absorbance values reached 0.6 to 0.8 at 600 nm. rHb expression was induced by the addition of 0.2 mM IPTG and the cultures were then supplemented with hemin (50 µg/ml) and glucose (20 g/L). Culture conditions and the protocol for preparing cell lysates were described previously (13, 14, 17-18). The overnight culture was saturated with CO for 15 min, and the cells were centrifuged for 45 min at 15000*g* and 4ºC. Pelleted cells were stored at -80ºC until further use. The pellets were resuspended with Tris lysis buffer (50 mM Tris, 0.5 mM DDT, 1mM EDTA) with lysozyme (1 mg/g of cells) for proper lysis before sonication. We used sonication cycles of 10 s pulse-on/20 s pulse-off for 15 min at 3.0 output (QSonicaQ500)(13, 17). We then added polyethyleneimine solution to the crude lysate to a final concentration of 0.25 to 1% to precipitate the bacterial nucleic acids. The crude lysate was then centrifuged for 45 min at 15000*g* and 4ºC, and the clarified supernatant was dialyzed overnight against the anion-exchange buffer for chromatography.

Recombinantly expressed proteins were purified using a two-step ion-exchange chromatography using the ÄKTA Start system (GE Healthcare) and Q-Sepharose columns (HiTrap QHP, 5 mL, 17-5159-01; GE Healthcare) pre-equilibrated with Tris/CAPS buffer. For each rHb, the buffer pH and other purification conditions were optimized on the basis of isoelectric point (pI)(Table S4). Each rHb was eluted using a linear gradient of 0-1.0 M NaCl. The eluted sample was desalted and dialyzed overnight against the second column buffer. We used prepackaged SP-Sepharose columns (HiTrap SPHP, 5 mL, 17-516101; GE Healthcare) equilibrated with HEPES/phosphate buffer (Table S4). Each rHb sample was eluted with a linear gradient of 0-1.0 M NaCl with the corresponding buffer and pH. The purified rHb samples were analyzed using 4-20% SDS-polyacrylamide gel electrophoresis to assess the purity of the fractions prior to *in vitro* measurements of $O_2$-binding properties.

**Collection of blood samples and preparation of hemolysates**. We sampled blood of alligator embryos at the 40% stage of pre-hatching development by catheterizing the major artery of the chorioallantoic membrane, as described by Bautista et al. (19). Upon completion of catheter placement, 100 to 500 μL of blood was collected into heparinized microhematocrit tubes and was then centrifuged at 20,854g for 5 min to separate red cell and plasma fractions. Individual hemolysates were prepared from thawed blood or erythrocyte samples (20-50 μL) by adding an approximately fourfold volume of ice-cold 10 mM HEPES, 0.5 mM EDTA, pH 7.4, followed by incubation on ice for 30 min. Lysed red blood cells were separated from the plasma fraction via centrifugation at 4°C (12,000 g, 15 min). Hemolysates were stored at -80°C prior to the isolation and purification of Hb fractions used for the functional experiments.

**Purification of native Hbs**. Thawed hemolysates were desalted on a PD-10 column (GE Healthcare) equilibrated with 20 nM HEPES, 0.5 nM EDTA, pH 7.65. To separate embryonic and adult isoHbs of American alligator (HbI and HbII, respectively), we subjected the hemolysates to anion-exchange chromatography on a Mono Q 5/50 GL column (1 mL) connected to an ÄKTA Pure Chromatography System (GE Healthcare). The column was equilibrated with 20 mM HEPES, 0.5 mM EDTA, pH 7.65, and isoHbs were eluted with a 0-0.1 M NaCl linear gradient at a flow rate of 1 mL/min. Absorbance was monitored at 415 and 280 nm to identify heme-containing proteins. Eluted peaks containing HbI and HbII were dialyzed against 10 mM HEPES, 0.5 mM EDTA, pH 7.4, and was then concentrated by ultrafiltration (Millipore) at 4°C to a final heme concentration of >0.05 mM.

**Measurement of oxygen equilibration curves**. Solutions of purified rHbs were desalted by passing samples through PD-10 desalting columns (GE Healthcare) that were equilibrated with 0.01 M HEPES, 0.5 mM EDTA, pH 7.4, and concentrated using Amicon Ultra-4 Centrifugal Filter Units (MilliporeSigma). $O_2$-equilibrium curves for Hb solutions (0.4 mM Hb tetramer in 100 mM HEPES, 0.5 mM EDTA buffer) were measured at 25°C using a Blood Oxygen Binding System (BOBS; Loligo® Systems). BOBS is a temperature-controlled gas diffusion chamber with a built-in spectrophotometer that allows the recording of changes in relative absorbance of a sample. Each Hb solution was sequentially equilibrated with 3-5 different partial pressures of $O_2$ ($PO_2$'s) at saturation levels between 30 to 70%, while absorbance was continually monitored at 430 nm (deoxy peak) and 421 nm (oxy/deoxy isosbestic point). Hb-$O_2$ saturation was estimated at each equilibrium step by comparing the absorbance at 430 nm to fully oxygenated and deoxygenated baselines. Estimates of the $PO_2$ at half-saturation ($P_{50}$) and the cooperativity coefficient ($n_{50}$) were then estimated from linear Hill plots (log[fractional saturation/[1-fractional saturation]] vs. log$P_{O2}$). $O_2$-equilibrium curves were measured in the absence (stripped) and presence of $Cl^-$ ions (0.1 M KCl) and organic phosphates (0.2 mM adenosine triphosphate [ATP]). $O_2$-equilibrium curves were also measured at three different pH levels, where the pH of working solutions was adjusted with NaOH to 7.2, 7.4, or 7.6 and were measured with an Orion Star A211 pH meter and an Orion™ PerpHecT™ ROSS™ Combination pH Micro Electrode. Linear regressions were fit to plots of log$P_{50}$ vs. pH, and the resulting equation was used to estimate $P_{50}$ values at pH 7.40 (± SE of the regression coefficient). The $n_{50}$ values are presented as mean ± SE of three measurements.

**Measurement of bicarbonate sensitivity**. Bicarbonate sensitivity of the ancestral and mutant rHbs was measured using 3 μL samples (0.14 mM heme, 0.1 M HEPES, pH 7.2, 0.5 mM EDTA at 25°C)(20-22). We measured time-course changes in $O_2$ saturation of the samples using BOBS. The system is connected to a programmable high precision Gas Mixing System pump (GMS – Loligo Systems) that allows for the control of discrete $O_2$ and $CO_2$ tensions – balanced with ultrapure $N_2$ – delivered to the surface of the sample in the chamber. Each rHb sample was equilibrated to the partial pressure of $O_2$ needed to reach its half-saturation ($P_{50}$), this value was estimated from the $O_2$ equilibrium curves described above. After equilibration to a $PO_2$ that matched the $P_{50}$ of the sample, 1% $CO_2$ was added to the gas mixture while maintaining a constant $PO_2$ in the chamber, and the reduction in Hb-$O_2$ saturation ($SO_2$) was recorded. Bicarbonate sensitivity was quantified as the log-transformed difference between $SO_2$ before and after the addition of 1% $CO_2$. The experiments were performed in triplicate and expressed as the mean ± 1 S.E.M.

**Tests of positive selection**. Using a maximum likelihood (ML) framework, we tested for evidence of positive selection in the crocodilian β-globin genes with codon-based models, as implemented in the *codeml* program in PAML v4.9 (23). We estimated phylogenetic relationships among crocodilian β-globin genes using IQTree version 2.1.3 (24) under the best-fitting model selected by the ModelFinder subroutine in IQ-Tree (25) and we used chicken $\beta^A$-globin as an outgroup (Fig. S6). We used the branch-site and clade models to examine variation in ω, the ratio of the rate of nonsynonymous substitution per nonsynonymous site, $d_N$, to the rate of synonymous substitution per synonymous site, $d_S$. Using both the branch and branch-site models (26, 27), we tested for changes in ω in the post-duplication line of descent leading from the single-copy progenitor of the *HBB-T1* and *HBB-T4* genes (β AncCrocT1/T4, which is also the common ancestor of all crocodilian β-type globin paralogs) to the most recent common ancestor of *HBB-T4* orthologs from all extant crocodilians (= the β-chain of AncCroc Hb)(Fig. S6). We examined (*i*) the branch connecting β AncCrocT1/T4 to the pre-duplication ancestor of *HBB-T4* and *HBB-T6*, (*ii*) the stem of the *HBB-T4* clade, (*iii*) the branch connecting β AncCrocT1/T4 to β AncCroc, and (*iv*) the entire *HBB-T4* clade, stem plus crown, relative to the rest of the tree. These 2-ω branch models were compared to a model where all branches share the same ω (M0). We followed a similar procedure for the branch-site models, setting the same 4 sets of branches as foreground. In background branches, sites evolve under purifying selection ($\omega_0$ <1) or under a neutral regime ($\omega_1$ =1), whereas in foreground branches, some sites are allowed to switch from purifying selection to positive selection ($\omega_{2a}$>1) or from a neutral regime to positive selection ($\omega_{2b}$>1). The null models for these analyses set $\omega_2$ =1.

**References – Supplementary Material**

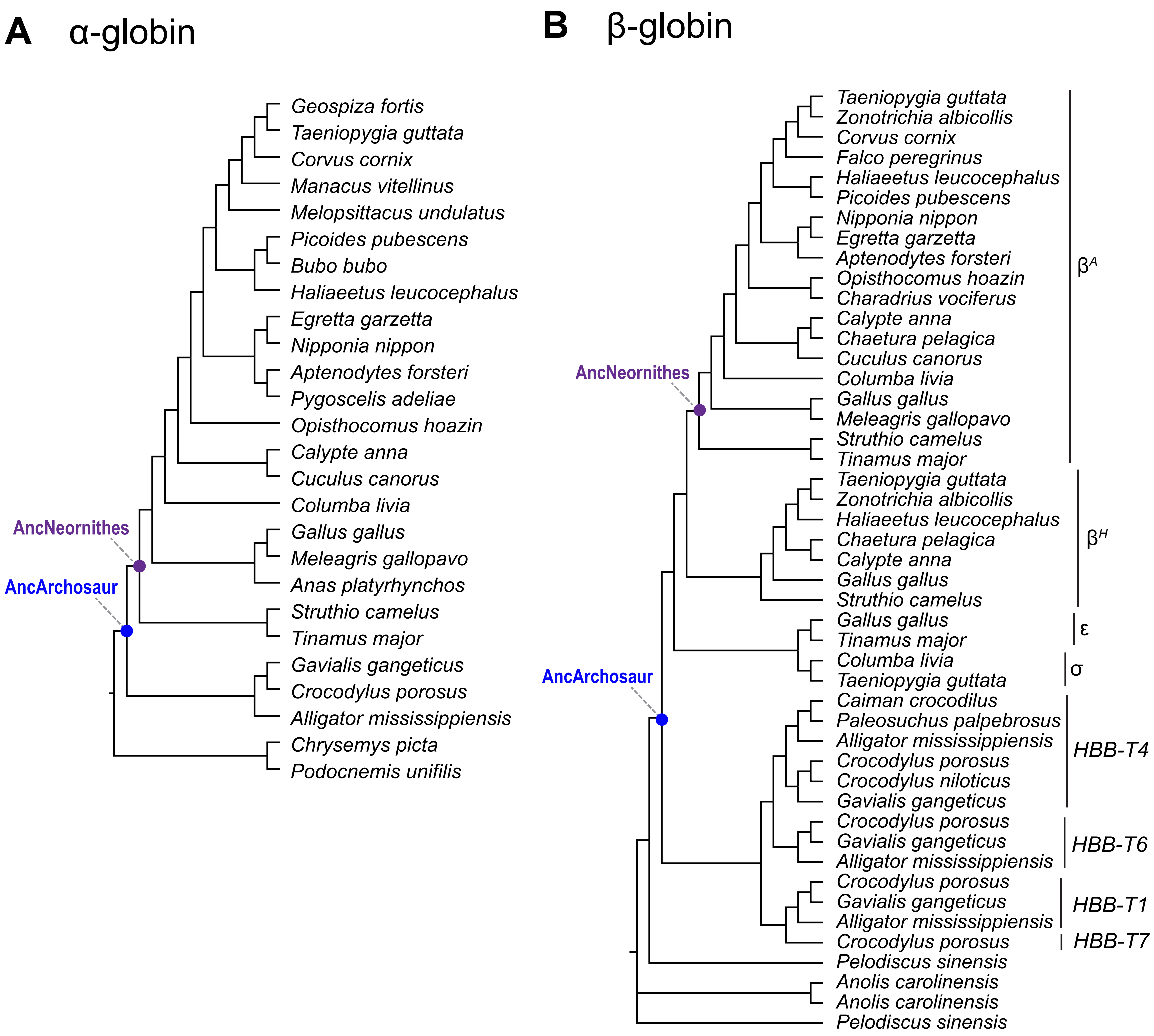


**Fig. S1. Phylogenetic trees of sauropsid globins used to estimate the ancestral α- and β-chain Hb sequences of the most recent common ancestors of archosaurs (AncArchosaur) and birds (AncNeornithes).** (A) Phylogeny of α-globin sequences and (B) β-globin sequences.

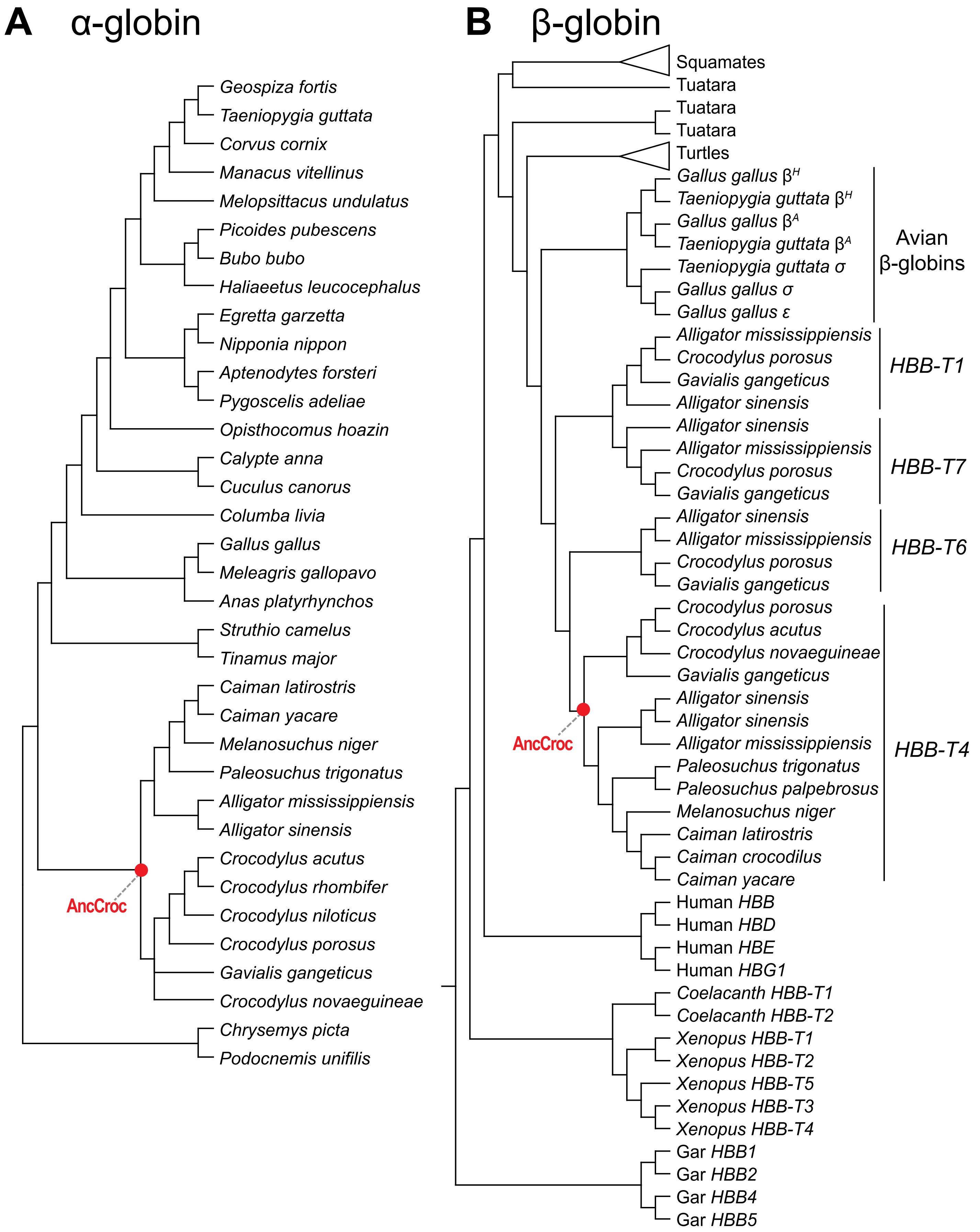


**Fig. S2. Phylogenetic trees of sauropsid globins used to estimate the ancestral α- and β-chain Hb sequences of the most recent common ancestor of crocodilians (AncCroc).** (A) Phylogeny of α-globin sequences and (B) β-globin sequences. The β-globin phylogeny includes representatives of all paralogous β-type globin genes that are products of crocodilian-specific duplication events (*HBB-T1*, *HBB-T4*, *HBB-T6*, and *HBB-T7*).

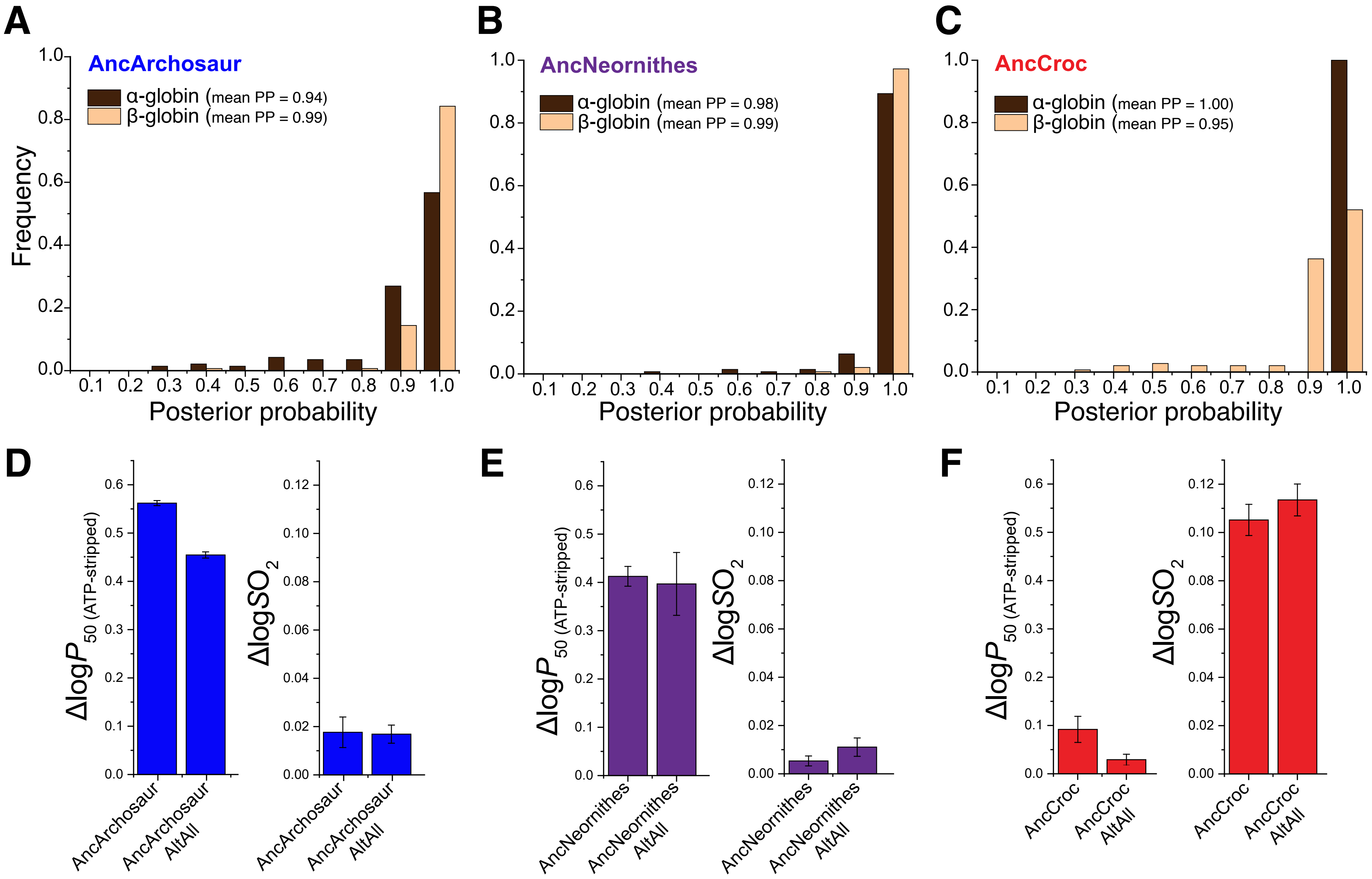


**Fig. S3. Ancestral sequence estimates and robustness of inferences about functional properties of ancestral proteins.** Distribution of site-specific posterior probabilities for estimated ancestral amino acid states in (A) AncArchosaur, (B) AncNeornithes, and (C) AncCroc. To assess the robustness of our functional inferences to uncertainties in ancestral sequence estimates, we synthesized additional rHbs that incorporated the alternative amino acid at each ambiguous site (posterior probability <0.80) in the corresponding ML sequence. In the case of (D) AncArchosaur, (E) AncNeornithes, and (F) AncCroc the ML ancestor and the corresponding AltAll protein have similar allosteric properties (sensitivity to ATP, insensitivity to bicarbonate in the case of AncArchosaur and AncNeornithes; sensitivity to bicarbonate, insensitivity to ATP in the case of AncCroc), indicating that inferences regarding evolved properties of the ancestral proteins are robust to statistical uncertainties in the ancestral sequence estimates.

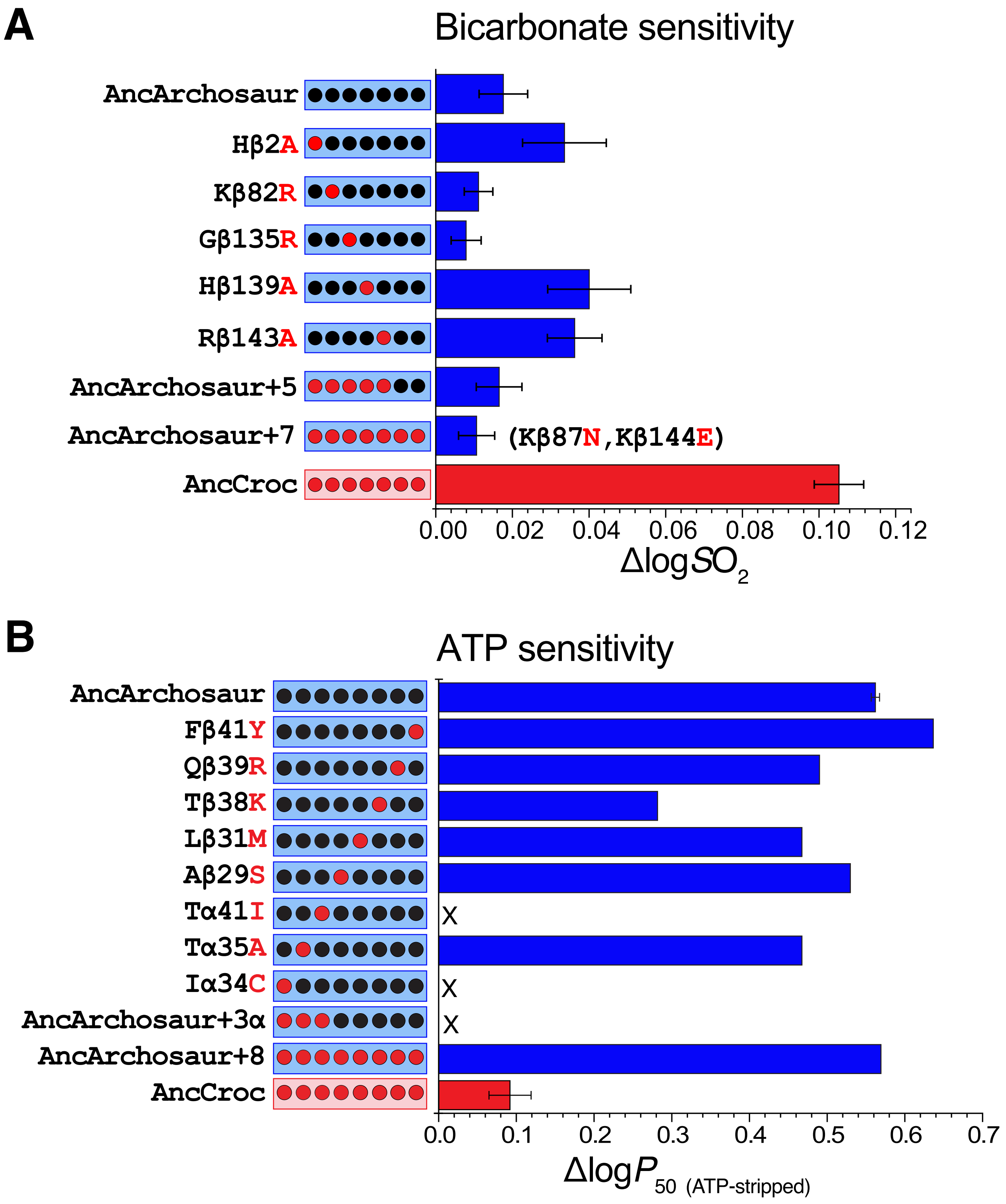


**Fig. S4. The gain and loss of different allosteric interactions involve distinct substitutions.** (A) The seven β-chain substitutions that were sufficient to eliminate ATP-sensitivity did not simultaneously enhance bicarbonate sensitivity on the AncArchosaur background. (B) Likewise, mutations at candidate sites for allosteric bicarbonate-binding did not significantly affect ATP-sensitivity on the AncArchosaur background.

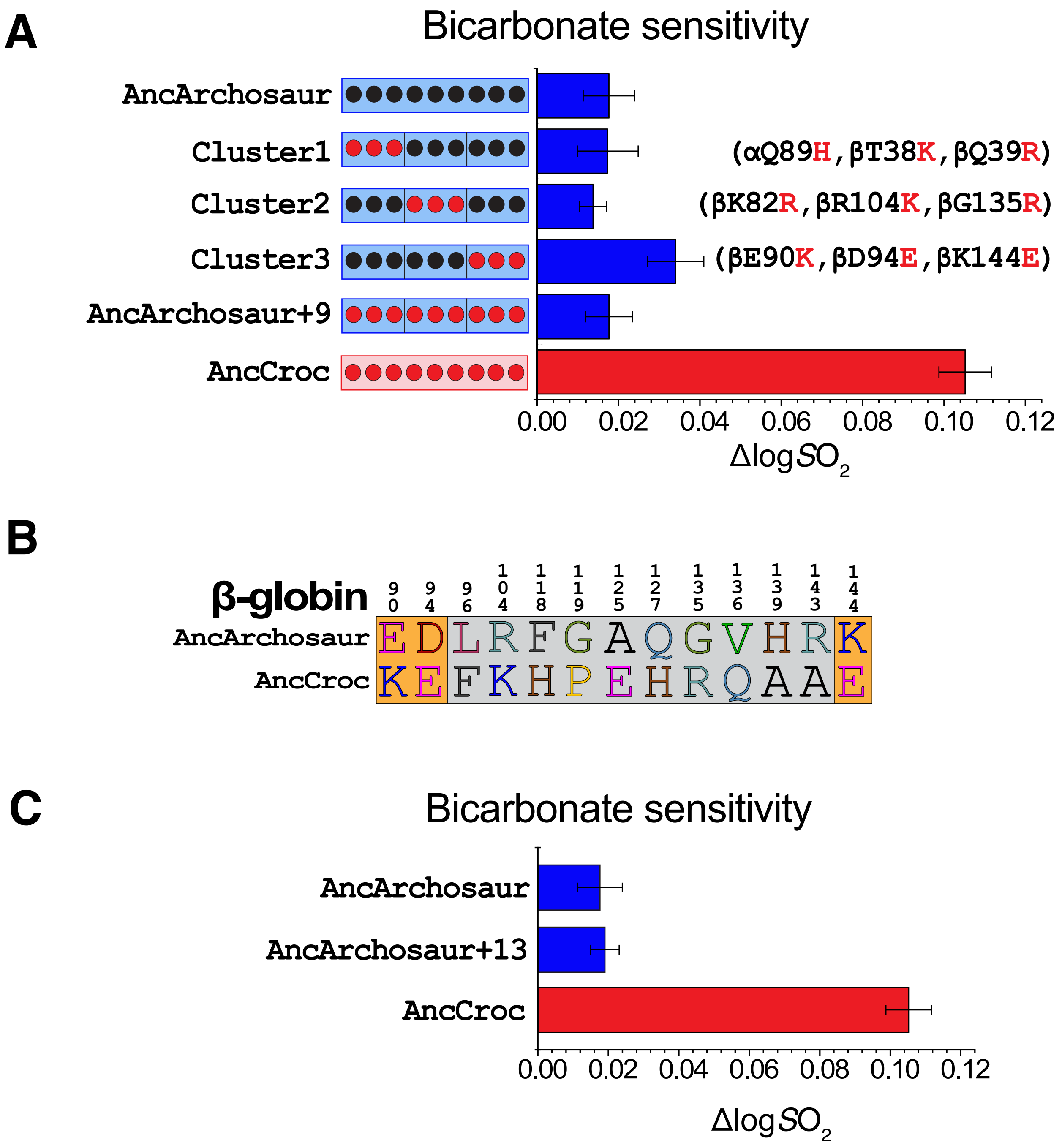


**Fig. S5. Testing effects of candidate charge-changing mutations on bicarbonate sensitivity.**
(A) Mutagenesis experiments tested 9 sites with binary combinations of ancestral and derived amino acids (black and red symbols, respectively). In the diagrams that show mutagenized sites, the AncArchosaur and AncCroc genetic backgrounds are denoted by blue and red, respectively. The 9 sites comprise 3 structurally discrete charge clusters, one of which (cluster 3) produced a modest increase in bicarbonate-sensitivity by itself, but not in combination with the remaining charge-changing mutations ('AncArchosaur+9'). (B) A follow-up experiment tested the 'cluster 3' charge-changing mutations (βE90K, βD94E, and βK144E; orange background) in combination with 10 additional croc-specific mutations at structurally proximal residues in the $\alpha_1\beta_2/\alpha_2\beta_1$ intersubunit interface (grey background). (C) The 13 mutations produced no increase in bicarbonate-sensitivity relative to the AncArchosaur reference value.

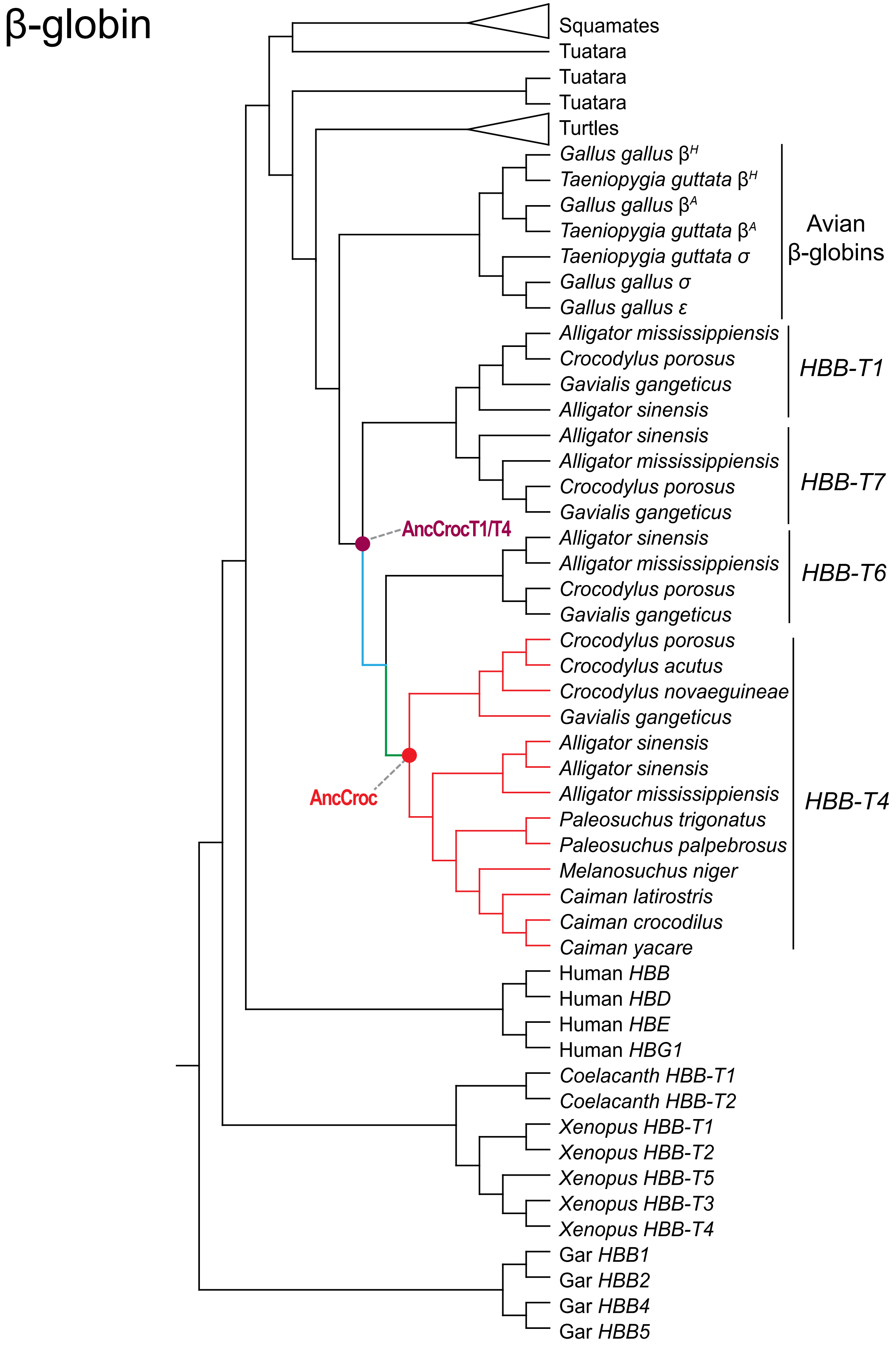


**Fig. S6. Phylogenetic tree of sauropsid β-globin genes used to test for evidence of positive selection in the β-chain of crocodilian Hb.** We tested for changes in ω (=dN/dS) in the post-duplication line of descent leading from the single-copy progenitor of the *HBB-T1* and *HBB-T4* genes (β AncCrocT1/T4, which is also the common ancestor of all crocodilian β-type globin paralogs) to the most recent common ancestor of adult-expressed *HBB-T4* orthologs from all extant crocodilians (= the β-chain of AncCroc Hb). We examined (i) the branch connecting β AncCrocT1/T4 to the pre-duplication ancestor of *HBB-T4* and *HBB-T6* (cyan branch), (ii) the stem of the *HBB-T4* clade (green branch), (iii) the branch connecting β AncCrocT1/T4 to β AncCroc (cyan branch + green branch), and (iv) the entire *HBB-T4* clade (green branch + red branches), stem plus crown, relative to the rest of the tree.

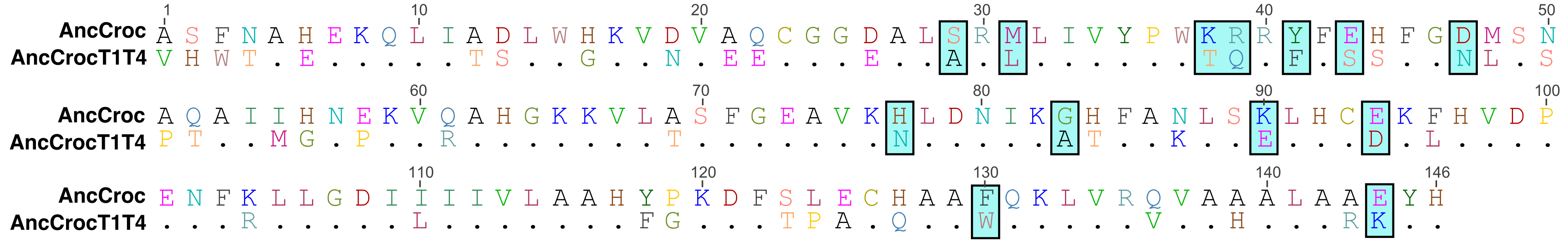


**Fig. S7. Sequences differences that distinguish the β-chain of AncCroc Hb from the single-copy progenitor of the HBB-T1 and HBB-T4 genes (β AncCrocT1/T4).** Boxes denote 13 sites where we introduced croc-specific mutations into the β AncCrocT1/T4 background: the eight β-chain mutations tested by Komiyama *et al*. in combination with five charge- or polarity-changing mutations concentrated in the α1β2/α2β1 interface.

**Table S1**. Oxygenation properties of all tested rHb mutants. $O_2$-affinity, $P_{50}$ (the partial pressure of $O_2$ at which Hb is half-saturated), and cooperativity of $O_2$-binding, $n_{50}$, were measured in the presence and absence of 0.2 mM ATP and 0.1 M KCl [25° C, pH 7.4]). Bicarbonate sensitivity was measured as the difference in log-transformed values of $SO_2$ [Hb-$O_2$ saturation] in the presence and absence of 1% $CO_2$. See *Supplementary Materials* for details. When introduced individually on the AncArchosaur background, two α-chain mutations, AncArc Iα34C and Tα41I (located in $\alpha_1\beta_1/\alpha_2\beta_2$ and $\alpha_1\beta_2/\alpha_2\beta_1$ interfaces, respectively), resulted in nonfunctional protein. The same was true when both mutations were combined with Tα35A as part of a trio of croc-specific α-chain mutations ('AncArchosaur+3α').

| Mutants | stripped | | +KCl | | +ATP | | KCl +ATP | | Δlog-$SO_2$ |
|---|---|---|---|---|---|---|---|---|---|
| | $P_{50}$ | $n_{50}$ | $P_{50}$ | $n_{50}$ | $P_{50}$ | $n_{50}$ | $P_{50}$ | $n_{50}$ | |
| AncNeornithes | 2.67±0.03 | 1.32±0.02 | 2.55±0.16 | 1.29±0.01 | 7.18±0.05 | 1.09±0.03 | 5.83±0.52 | 1.20±0.03 | 0.0053±0.0021 |
| AncNeornithes AltAll | 2.31±0.32 | 1.30±0.02 | 2.49±0.09 | 1.30±0.02 | 5.85±0.10 | 1.11±0.02 | 4.32±0.30 | 1.18±0.04 | 0.0111±0.0038 |
| AncArchosaur | 2.20±0.06 | 1.41±0.04 | 2.40±0.07 | 1.65±0.03 | 7.96±0.13 | 1.29±0.06 | 6.54±0.46 | 1.44±0.11 | 0.0176±0.0063 |
| AncArchosaur AltAll | 2.20±0.06 | 1.51±0.05 | 2.29±0.24 | 1.71±0.06 | 6.27±0.20 | 1.73±0.00 | 5.50±0.19 | 1.37±0.15 | 0.0169±0.0037 |
| AncCroc | 2.92±0.16 | 1.01±0.02 | 3.46±0.06 | 0.91±0.01 | 3.75±0.70 | 0.93±0.03 | 3.31±0.13 | 0.97±0.03 | 0.1052±0.0065 |
| AncCroc AltAll | 2.07±0.02 | 1.04±0.01 | 3.01±0.21 | 1.23±0.01 | 2.22±0.04 | 1.10±0.00 | 3.09±0.10 | 1.22±0.03 | 0.1135±0.0066 |
| AncArc Hβ2A | 2.46±0.09 | 1.60±0.06 | 2.70±0.12 | 1.64±0.04 | 8.81±0.23 | 1.62±0.02 | 7.25±0.17 | 1.63±0.02 | 0.0336±0.0109 |
| AncArc Kβ82R | 2.41±0.20 | 1.40±0.02 | 2.45±0.07 | 1.42±0.01 | 5.12±0.25 | 1.56±0.05 | 3.64±0.13 | 1.51±0.09 | 0.0112±0.0037 |
| AncArc Gβ135R | 2.34±0.09 | 1.38±0.04 | 2.61±0.14 | 1.39±0.07 | 5.55±0.15 | 1.48±0.07 | 4.79±0.38 | 1.54±0.02 | 0.0079±0.0039 |
| AncArc Hβ139R | 2.28±0.06 | 1.48±0.11 | 2.52±0.00 | 1.42±0.03 | 6.07±0.34 | 1.23±0.02 | 4.25±0.02 | 1.65±0.04 | 0.0400±0.0109 |
| AncArc Rβ143A | 5.03±0.01 | 1.63±0.01 | 4.92±0.08 | 1.76±0.01 | 10.69±0.29 | 1.79±0.14 | 6.97±0.26 | 1.99±0.01 | 0.0362±0.0071 |
| AncArchosaur+5 | 2.48±0.25 | 1.43±0.01 | 2.85±0.07 | 1.56±0.04 | 4.51±0.24 | 1.74±0.12 | 3.27±0.14 | 1.69±0.07 | 0.0165±0.0060 |
| AncArchosaur+7 | 3.70±0.03 | 1.50±0.02 | 4.88±0.31 | 1.64±0.05 | 4.41±0.11 | 1.65±0.02 | 5.07±0.13 | 1.61±0.03 | 0.0107±0.0047 |
| AncArc Tα35A | 1.21±0.04 | 1.02±0.05 | - | - | 3.55±0.05 | 0.89±0.02 | - | - | 0.0030 |
| AncArc Aβ29S | 0.88±0.02 | 1.13±0.05 | - | - | 2.98±0.09 | 0.77±0.03 | - | - | 0.0046 |
| AncArc Lβ31M | 0.91±0.03 | 0.96±0.07 | - | - | 2.67±0.06 | 0.92±0.03 | - | - | 0.0219±0.0019 |
| AncArc Tβ38K | 1.15±0.01 | 1.03±0.03 | - | - | 2.20±0.04 | 0.91±0.03 | - | - | 0.0113±0.0045 |
| AncArc Qβ39R | 1.43±0.03 | 1.19±0.05 | - | - | 4.42±0.11 | 0.93±0.03 | - | - | 0.0017±0.0025 |
| AncArc Fβ41Y | 1.60±0.04 | 1.12±0.07 | - | - | 6.93±0.16 | 0.76±0.03 | - | - | 0.0113 |
| AncArc Tβ38K_Fβ41Y | 1.17±0.03 | 0.60±0.02 | - | - | 2.61±0.05 | 0.42±0.02 | - | - | 0.0106±0.0021 |
| AncArchosaur+8 | 0.89±0.02 | 0.97±0.04 | - | - | 2.37±0.11 | 0.82±0.06 | - | - | 0.0237±0.0048 |
| AncArchosaur+18 | 7.49±0.15 | 1.41±0.16 | 7.79±0.32 | 1.27±0.02 | 7.40±0.11 | 1.29±0.02 | 6.85±0.92 | 1.39±0.02 | 0.0620±0.0022 |
| AncArchosaur cluster1 | 2.23±0.20 | 1.23±0.12 | 2.12±0.06 | 1.23±0.05 | 2.75±0.27 | 1.12±0.03 | 2.31±0.14 | 1.16±0.02 | 0.0173±0.0075 |

| | | | | | | | | | |
|---|---|---|---|---|---|---|---|---|---|
| AncArchosaur cluster2 | 2.35±0.33 | 1.26±0.03 | 2.50±0.29 | 1.31±0.04 | 3.88±0.22 | 1.22±0.05 | 3.35±0.16 | 1.27±0.08 | 0.0137±0.0034 |
| AncArchosaur cluster3 | 6.88±0.47 | 1.51±0.08 | 6.21±0.45 | 1.50±0.04 | 16.96±1.44 | 1.78±0.12 | 11.85±1.02 | 1.81±0.07 | 0.0340±0.0069 |
| AncArchosaur+9 | 4.61±0.12 | 1.09±0.02 | 4.33±0.20 | 1.07±0.03 | 5.27±0.22 | 1.18±0.02 | 4.90±0.49 | 1.15±0.03 | 0.0177±0.0058 |
| AncArchosaur +13 | 1.36±0.13 | 1.02±0.02 | 1.44±0.02 | 1.01±0.01 | 2.10±0.17 | 0.94±0.04 | 1.94±0.15 | 0.97±0.05 | 0.0190±0.0040 |
| AncCrocT1T4 | 4.12±0.06 | 1.38±0.01 | 4.94±0.55 | 1.37±0.02 | 5.32±0.14 | 1.32±0.02 | 4.92±0.09 | 1.33±0.04 | 0.0079±0.0018 |
| AncCrocT1T4 AltAll | 2.16±0.08 | 1.07±0.03 | 2.54±0.12 | 1.04±0.02 | 4.64±0.46 | 0.98±0.02 | 4.03±0.21 | 1.02±0.01 | 0.0257±0.0059 |
| AncCrocT1T4+13 | 4.31±0.13 | 0.97±0.01 | 4.32±0.07 | 0.95±0.01 | 6.09±0.32 | 0.98±0.03 | 6.93±0.19 | 0.96±0.03 | 0.0640±0.0150 |
| AncCroc -13 | 1.79±0.11 | 0.95±0.07 | 2.08±0.05 | 0.89±0.08 | 3.01±0.19 | 1.21±0.14 | 1.82±0.05 | 0.90±0.00 | 0.0157±0.0055 |
| AncCrocT1T4+5 | 2.99±0.02 | 1.00±0.02 | 3.21±0.10 | 1.04±0.02 | 5.77±0.05 | 0.90±0.02 | 6.01±0.11 | 0.91±0.02 | 0.0575±0.0068 |
| AncCroc -5 | 4.03±0.31 | 0.97±0.12 | 3.06±0.16 | 0.62±0.11 | 3.23±0.48 | 0.74±0.10 | 3.59±0.13 | 0.89±0.14 | 0.0172±0.0046 |
| AncCrocT1T4-Part I | 2.16±0.06 | 1.08±0.03 | 2.53±0.09 | 1.15±0.02 | 4.66±0.17 | 1.18±0.02 | 4.34±0.20 | 1.10±0.05 | 0.1049±0.0063 |
| AncCrocT1T4-Part II | 4.41±0.02 | 1.10±0.01 | 3.14±0.08 | 0.98±0.02 | 5.32±0.22 | 0.99±0.05 | 6.47±0.29 | 1.03±0.02 | 0.0173±0.0062 |
| AncCrocT1T4-Part III | 2.68±0.03 | 1.05±0.03 | 1.85±0.11 | 0.91±0.04 | 2.30±0.08 | 1.01±0.02 | 2.09±0.04 | 1.00±0.02 | 0.0182±0.0025 |
| AncCrocT1T4-Part IA | 2.56±0.00 | 0.94±0.00 | 2.56±0.18 | 0.89±0.03 | 3.98±0.16 | 0.84±0.04 | 3.22±0.14 | 0.86±0.01 | 0.0220±0.0030 |
| AncCrocT1T4-Part IB | 1.81±0.03 | 0.89±0.04 | 1.76±0.32 | 0.75±0.03 | 2.80±0.27 | 0.86±0.05 | 1.98±0.09 | 0.85±0.02 | 0.0876±0.0073 |
| AncCrocT1T4- Part IB1 | 2.77±0.35 | 0.96±0.05 | 3.03±0.07 | 0.99±0.01 | 2.11±0.19 | 0.73±0.04 | 3.39±0.20 | 0.97±0.02 | 0.0738±0.0082 |
| Human Hb | 9.05±0.74 | 2.11±0.10 | 17.74±0.46 | 2.15±0.05 | 10.25±0.58 | 2.14±0.05 | 17.40±0.18 | 2.02±0.06 | 0.0173±0.0036 |
| Hb Sobek | 3.20±0.01 | 1.52±0.03 | 5.08±0.86 | 1.20±0.09 | 3.26±0.06 | 1.35±0.04 | 6.24±0.34 | 1.12±0.03 | 0.0255±0.0015 |

**Table S2**. Branch model statistics for crocodilian β-globin paralogs. ω= $d_N/d_S$ (the rate of nonsynonymous substitutions per nonsynonymous site relative to the rate of synonymous substitutions per synonymous site). Models considered the following branches as foreground: (*i*) the branch connecting β AncCrocT1/T4 to the pre-duplication ancestor of *HBB-T4* and *HBB-T6*, (*ii*) the branch connecting β AncCrocT1/T4 to *HBB-T4* (= β AncCroc), (*iii*) the stem of the *HBB-T4* clade, and (*iv*) the entire *HBB-T4* clade, stem plus crown, relative to the rest of the tree (see fig. S6). NS = Not significant.

| **Model** | **np** | **-lnL** | **ω estimates** | **LRT** |
|---|---|---|---|---|
| Model 0 (null model) | 61 | 2887.22 | ω = 0.2245 | |
| AncCrocT1/T4 to AncCrocT4/T6 branch as foreground | 62 | 2884.14 | $\omega_0 = 0.21$<br>$\omega_1 = 2.99$ | $P < 0.01$ |
| AncCrocT1/T4 to β AncCroc as foreground | 62 | 2887.21 | $\omega_0 = 0.22$<br>$\omega_1 = 0.23$ | NS |
| Stem of *HBB-T4* as foreground | 62 | 2887.22 | $\omega_0 = 0.24$<br>$\omega_1 = 0.14$ | NS |
| *HBB-T4* clade as foreground (stem + crown) | 62 | 2872.1 | $\omega_0 = 0.14$<br>$\omega_1 = 0.50$ | $P < 0.001$ |

**Table S3**. Branch-site model statistics for reconstructed ancestral crocodilian β-globin genes. NS = Not significant.

| **Model** | **np** | **-lnL** | **Site Class** | **%** | **$\omega_0$** | **$\omega_1$** | **LRT** |
|---|---|---|---|---|---|---|---|
| AncCrocT1/T4 to AncCrocT4/T6 as foreground | | | | | | | |
| Model A null | 63 | 2862.12 | 0 | 0.55 | 0.16 | 0.16 | $P < 0.01$ |
| | | | 1 | 0.12 | 1.00 | 1.00 | |
| | | | 2a | 0.27 | 0.16 | 1.00 | |
| | | | 2b | 0.06 | 1.00 | 1.00 | |
| Model A | 64 | 2857.50 | 0 | 0.74 | 0.16 | 0.16 | |
| | | | 1 | 0.10 | 1.00 | 1.00 | |
| | | | 2a | 0.10 | 0.16 | 30.59 | |
| | | | 2b | 0.02 | 1.00 | 30.59 | |
| AncCrocT1/T4 to *HBB-T4* (= βAncCroc) as foreground | | | | | | | |
| Model A null | 63 | 2860.40 | 0 | 0.69 | 0.14 | 0.14 | $P < 0.01$ |
| | | | 1 | 0.15 | 1.00 | 1.00 | |
| | | | 2a | 0.14 | 0.14 | 1.00 | |
| | | | 2b | 0.03 | 1.00 | 1.00 | |
| Model A | 64 | 2856.50 | 0 | 0.74 | 0.15 | 0.15 | |
| | | | 1 | 0.14 | 1.00 | 1.00 | |
| | | | 2a | 0.10 | 0.15 | 5.55 | |
| | | | 2b | 0.02 | 1.00 | 5.55 | |
| Stem of *HBB-T4* clade as foreground | | | | | | | |
| Model A null | 63 | 2862.12 | 0 | 0.70 | 0.14 | 0.14 | *NS* |
| | | | 1 | 0.20 | 1.00 | 1.00 | |
| | | | 2a | 0.08 | 0.14 | 1.00 | |
| | | | 2b | 0.02 | 1.00 | 1.00 | |
| Model A | 64 | 2858.02 | 0 | 0.70 | 0.15 | 0.15 | |
| | | | 1 | 0.20 | 1.00 | 1.00 | |
| | | | 2a | 0.07 | 0.15 | 19.20 | |
| | | | 2b | 0.02 | 1.00 | 19.20 | |

| | | | | | | | |
|---|---|---|---|---|---|---|---|
| *HBB-T4* clade as foreground (stem + crown) | | | | | | | |
| Model A null | 63 | 2835.73 | 0 | 0.58 | 0.10 | 0.10 | |
| | | | 1 | 0.08 | 1.00 | 1.00 | |
| | | | 2a | 0.30 | 0.10 | 1.00 | |
| | | | 2b | 0.04 | 1.00 | 1.00 | *P* ~ 0.0001 |
| Model A | 64 | 2830.69 | 0 | 0.62 | 0.11 | 0.11 | |
| | | | 1 | 0.09 | 1.00 | 1.00 | |
| | | | 2a | 0.26 | 0.11 | 2.01 | |
| | | | 2b | 0.04 | 1.00 | 2.01 | |

**Table S4**. Buffer conditions for purification of recombinantly expressed Hb mutants via two-step ion-exchange chromatography.

| Mutants | Q-column Buffer | SP-column Buffer |
|---|---|---|
| AncNeornithes | 20 mM Tris, 0.5mM EDTA, 1 mM DTT pH 8.4 | 20 mM HEPES with 0.5mM EDTA, 1 mM DTT pH 7.0 |
| AncNeornithes AltAll | 20 mM Tris, 0.5mM EDTA, 1 mM DTT pH 8.4 | 20 mM HEPES with 0.5mM EDTA, 1 mM DTT pH 7.0 |
| AncArchosaur | 20 mM Tris, 0.5mM EDTA, 1 mM DTT pH 8.8 | 20 mM HEPES with 0.5mM EDTA, 1 mM DTT pH 7.0 |
| AncArchosaur AltAll | 20 mM Tris, 0.5mM EDTA, 1 mM DTT pH 8.8 | 20 mM HEPES with 0.5mM EDTA, 1 mM DTT pH 7.0 |
| AncCroc | 20 mM Tris, 0.5mM EDTA, 1 mM DTT pH 8.8 | 50 mM Phosphate, 0.5 mM EDTA,0.5 mM IHP pH 6.2 |
| AncCroc AltAll | 20 mM CAPS, 0.5mM EDTA, 1 mM DTT pH 9.4 | 50 mM Phosphate, 0.5 mM EDTA,0.5 mM IHP pH 6.4 |
| AncArc Hβ2A | 20 mM Tris, 0.5mM EDTA, 1 mM DTT pH 8.8 | 20 mM HEPES with 0.5mM EDTA, 1 mM DTT pH 7.0 |
| AncArc Kβ82R | 20 mM Tris, 0.5mM EDTA, 1 mM DTT pH 8.8 | 20 mM HEPES with 0.5mM EDTA, 1 mM DTT pH 7.0 |
| AncArc Gβ135R | 20 mM Tris, 0.5mM EDTA, 1 mM DTT pH 8.8 | 20 mM HEPES with 0.5mM EDTA, 1 mM DTT pH 7.0 |
| AncArc Hβ139R | 20 mM Tris, 0.5mM EDTA, 1 mM DTT pH 8.8 | 20 mM HEPES with 0.5mM EDTA, 1 mM DTT pH 7.0 |
| AncArc Rβ143A | 20 mM Tris, 0.5mM EDTA, 1 mM DTT pH 8.8 | 20 mM HEPES with 0.5mM EDTA, 1 mM DTT pH 7.0 |
| AncArchosaur+5 | 20 mM Tris, 0.5mM EDTA, 1 mM DTT pH 8.8 | 20 mM HEPES with 0.5mM EDTA, 1 mM DTT pH 7.0 |
| AncArchosaur+7 | 20 mM Tris, 0.5mM EDTA, 1 mM DTT pH 8.8 | 20 mM HEPES with 0.5mM EDTA, 1 mM DTT pH 7.0 |
| AncArc Tα35A | 20 mM Tris, 0.5mM EDTA, 1 mM DTT pH 8.8 | 20 mM HEPES with 0.5mM EDTA, 1 mM DTT pH 7.0 |
| AncArc Aβ29S | 20 mM Tris, 0.5mM EDTA, 1 mM DTT pH 8.8 | 20 mM HEPES with 0.5mM EDTA, 1 mM DTT pH 7.0 |
| AncArc Lβ31M | 20 mM Tris, 0.5mM EDTA, 1 mM DTT pH 8.8 | 20 mM HEPES with 0.5mM EDTA, 1 mM DTT pH 7.0 |
| AncArc Tβ38K | 20 mM Tris, 0.5mM EDTA, 1 mM DTT pH 8.8 | 20 mM HEPES with 0.5mM EDTA, 1 mM DTT pH 7.0 |
| AncArc Qβ39R | 20 mM Tris, 0.5mM EDTA, 1 mM DTT pH 8.8 | 20 mM HEPES with 0.5mM EDTA, 1 mM DTT pH 7.0 |
| AncArc Fβ41Y | 20 mM Tris, 0.5mM EDTA, 1 mM DTT pH 8.8 | 20 mM HEPES with 0.5mM EDTA, 1 mM DTT pH 7.0 |
| AncArc Tβ38K_Fβ41Y | 20 mM Tris, 0.5mM EDTA, 1 mM DTT pH 8.8 | 20 mM HEPES with 0.5mM EDTA, 1 mM DTT pH 7.0 |
| AncArchosaur+8 | 20 mM Tris, 0.5mM EDTA, 1 mM DTT, pH 8.4 | 20 mM HEPES with 0.5mM EDTA, 1 mM DTT pH 7.0 |
| AncArchosaur+18 | 20 mM CAPS, 0.5mM EDTA, 1 mM DTT, pH 9.3 | 50 mM Phosphate, 0.5 mM EDTA pH 7.0 |
| AncArchosaur cluster1 | 20 mM Tris, 0.5mM EDTA, 1 mM DTT pH 8.8 | 20 mM HEPES with 0.5mM EDTA, 1 mM DTT pH 7.0 |
| AncArchosaur cluster2 | 20 mM Tris, 0.5mM EDTA, 1 mM DTT pH 8.8 | 20 mM HEPES with 0.5mM EDTA, 1 mM DTT pH 7.0 |
| AncArchosaur cluster3 | 20 mM Tris, 0.5mM EDTA, 1 mM DTT pH 8.8 | 20 mM HEPES with 0.5mM EDTA, 1 mM DTT pH 6.8 |
| AncArchosaur+9 | 20 mM Tris, 0.5mM EDTA, 1 mM DTT pH 8.8 | 20 mM HEPES with 0.5mM EDTA, 1 mM DTT pH 7.0 |
| AncArchosaur +13 | 20 mM Tris, 0.5mM EDTA, 1 mM DTT pH 8.8 | 20 mM HEPES with 0.5mM EDTA, 1 mM DTT pH 6.8 |
| AncCrocT1T4 | 20 mM Tris, 0.5mM EDTA, 1 mM DTT pH 9.5 | 50 mM Phosphate, 0.5 mM EDTA, pH 6.2 |
| AncCrocT1T4 AltAll | 20 mM Tris, 0.5mM EDTA, 1 mM DTT pH 9.5 | 50 mM Phosphate, 0.5 mM EDTA, pH 6.2 |

| | | |
|---|---|---|
| AncCrocT1T4+13 | 20 mM Tris, 0.5mM EDTA, 1 mM DTT pH 9.5 | 50 mM Phosphate, 0.5 mM EDTA, pH 6.2 |
| AncCroc -13 | 20 mM Tris, 0.5mM EDTA, 1 mM DTT pH 8.8 | 50 mM Phosphate, 0.5 mM EDTA, pH 6.2 |
| AncCrocT1T4+5 | 20 mM Tris, 0.5mM EDTA, 1 mM DTT pH 9.5 | 50 mM Phosphate, 0.5 mM EDTA, pH 6.2 |
| AncCroc -5 | 20 mM Tris, 0.5mM EDTA, 1 mM DTT pH 8.8 | 50 mM Phosphate, 0.5 mM EDTA, pH 6.2 |
| AncCrocT1T4-Part I | 20 mM Tris, 0.5mM EDTA, 1 mM DTT pH 8.8 | 50 mM Phosphate, 0.5 mM EDTA, pH 6.2 |
| AncCrocT1T4-Part II | 20 mM Tris, 0.5mM EDTA, 1 mM DTT pH 8.8 | 50 mM Phosphate, 0.5 mM EDTA, pH 6.2 |
| AncCrocT1T4-Part III | 20 mM Tris, 0.5mM EDTA, 1 mM DTT pH 8.8 | 50 mM Phosphate, 0.5 mM EDTA, pH 6.2 |
| AncCrocT1T4-Part IA | 20 mM Tris, 0.5mM EDTA, 1 mM DTT pH 8.8 | 50 mM Phosphate, 0.5 mM EDTA, pH 6.2 |
| AncCrocT1T4-Part IB | 20 mM Tris, 0.5mM EDTA, 1 mM DTT pH 8.8 | 50 mM Phosphate, 0.5 mM EDTA, pH 6.2 |
| AncCrocT1T4- Part IB1 | 20 mM Tris, 0.5mM EDTA, 1 mM DTT pH 8.8 | 50 mM Phosphate, 0.5 mM EDTA, pH 6.2 |
| Human Hb | 20 mM Tris, 0.5mM EDTA, pH 8.8 | 20 mM HEPES with 0.5 mM EDTA, pH 7.0 |
| Hb Sobek | 20 mM Tris, 0.5mM EDTA, pH 8.8 | 20 mM HEPES with 0.5 mM EDTA, pH 7.0 |
